# *In vitro* comparison of Aβ-targeting SNIPR, synNotch, and TRUCK for cell-based drug delivery in Alzheimer’s disease

**DOI:** 10.64898/2026.04.29.721717

**Authors:** Cynthia J. Siebrand, Zachary Mayeri, Ian Brown, Julie K. Andersen, Chaska C. Walton

## Abstract

Pioneering research is adapting chimeric antigen receptors (CARs) from oncology to Alzheimer’s disease (AD) by targeting amyloid beta (Aβ). Newer synthetic receptor systems can go beyond, transforming cells into targeted biological drug factories that can couple Aβ detection to synthesis and secretion of genetically encoded therapeutics. Among candidate systems, T cells Redirected for Universal Cytokine Killing (TRUCK), synthetic Notch (synNotch), and Synthetic Intramembrane Proteolysis Receptors (SNIPR) have shown promise in oncology. Here, we adapt these platforms to AD using a shared Aβ-targeting binding domain derived from Aducanumab (Aduhelm), coupled to inducible expression cassettes driving identical transgenes: secreted Metridia luciferase (MetLuc) and a Lecanemab (Leqembi)-based chimeric human-mouse antibody (chLecanemab). To validate these systems *in vitro*, Jurkat clones expressing each receptor were treated with oligomer-enriched Aβ (AβO) to model AD, and receptor output was quantified by media MetLuc levels and chLecanemab colocalization with Aβ aggregates. For TRUCK systems, we show the Aβ-targeting CAR successfully activated Jurkat cells by flow cytometry. We also show that six Nuclear Factor of Activated T-cells (NFAT) tandem repeat response elements (6xNFAT) paired with either minimal interleukin-2, synthetic TATA box, or minimal cytomegalovirus promoters resulted in functional regulatory regions. Despite this, all TRUCK variants failed to significantly upregulate MetLuc in response to AβO. In contrast, both synNotch and SNIPR responded robustly to AβO, with SNIPR outperforming synNotch in both MetLuc and chLecanemab production. These findings establish SNIPR and synNotch as promising platforms for future research on cell-based targeted therapeutic delivery in AD.

## Introduction

AD poses a major challenge to an aging society, with FDA-approved Aβ-clearing antibodies currently serving as our primary line of defense (Cummings *et al*., 2024). However, existing treatments are insufficient to fully halt disease progression, spurring extensive research into new strategies and targets for clinical trials (Cummings *et al*., 2025). One radical new approach in preclinical research involves adapting the synthetic biology principles underlying CAR T cell (CAR-T) cancer immunotherapy to AD (Saetzler *et al*., 2023; Siebrand *et al*., 2025; Boskovic *et al*., 2026; Kim *et al*., 2024; Chen *et al*., 2026; Heiss *et al*., 2025; Bergo *et al*., 2025; Spetz *et al*., 2025).

In CAR-T therapy for oncology, a CAR enables engineered immune cells to recognize specific membrane-bound ligands on cancer cells to redirect their cytotoxicity toward malignancies (June and Sadelain, 2018). CAR-based strategies in neurodegenerative diseases such as AD differ in several respects from CAR-T therapy in cancer. First, CAR technologies are being adapted to target extracellular protein aggregates rather than ligands on the cell membrane (Saetzler *et al*., 2023; Kim *et al*., 2024; Heiss *et al*., 2025; Siebrand *et al*., 2025; Boskovic *et al*., 2026; Chen *et al*., 2026). Second, the objective is not to kill cells but to activate non-cytotoxic immunomodulatory functions in the vicinity of the targeted extracellular protein aggregates. Consequently, CARs and modified CARs explored for AD are generally expressed in non-cytotoxic cells.

As one of the defining neuropathological hallmarks of AD (Hyman *et al*., 2012; Jack *et al*., 2018; DeTure and Dickson, 2019), extracellular Aβ deposits constitute a highly attractive homing target for engineered cells (Saetzler *et al*., 2023; Kim *et al*., 2024; Heiss *et al*., 2025; Siebrand *et al*., 2025; Boskovic *et al*., 2026; Chen *et al*., 2026). Aβ is generated from amyloid precursor protein (APP) through sequential cleavage by β- and γ-secretase, and can assemble into extracellular oligomers, protofibrils, and fibrils, contributing to amyloid plaque formation (Hampel *et al*., 2021). Our previous work validated multiple CARs targeting different extracellular Aβ preparations, including pyroglutamated Aβ associated with parenchymal cored plaques, as well as a distinct CAR targeting tau preformed fibrils (Siebrand *et al*., 2025). Others have shown that Aβ-targeting CAR CD4⁺ T cells significantly reduced amyloid burden and reactive glial responses in the 5xFAD mouse model (Boskovic *et al*., 2026). Additionally, Aβ-targeting CAR-engineered macrophages (CAR-M) effectively cleared Aβ plaques in APP/PS1 mice (Kim *et al*., 2024), while Aβ-targeting CAR-engineered astrocytes (CAR-A) similarly reduced amyloid burden in a 5xFAD mouse model (Chen *et al*., 2026).

Conventional CAR-based interventions primarily leverage the endogenous effector functions of the engineered cell. For example, an Aβ-targeting CAR in regulatory T cells (Tregs) elicited immunosuppression *in vitro* (Saetzler *et al*., 2023), whereas in macrophages, microglia, and astrocytes, CAR technology induced phagocytic activity (Kim *et al*., 2024; Heiss *et al*., 2025; Chen *et al*., 2026). Thus, CAR applications for AD are largely constrained by the natural effector functions of the cell type in which it is deployed. However, beyond CARs, new synthetic receptor systems can be used to bestow cells with entirely new artificial functions (Manhas *et al*., 2022). Specifically, cells can be reprogrammed to act as biological drug factories, capable of sensing pathological signals and responding by synthesizing and secreting therapeutic biologics on demand. Arguably, the most relevant of these synthetic receptor systems are TRUCK, synNotch, and SNIPRs (Chmielewski *et al*., 2011; Chmielewski and Abken, 2020; Zhu *et al*., 2022; Manhas *et al*., 2022; Morsut *et al*., 2016; Roybal *et al*., 2016). These systems link antigen recognition to inducible transgene expression, enabling controlled production of protein payloads such as antibodies and cytokines.

TRUCK systems, also referred to as fourth-generation CARs (Chmielewski and Abken, 2015), were originally developed in oncology to enhance CAR-mediated antigen-specific effector functions by adding inducible local cytokine release within the tumor microenvironment (Chmielewski *et al*., 2011; Chmielewski and Abken, 2017, 2020; Kunert *et al*., 2018; Liu *et al*., 2019; Zimmermann *et al*., 2020; Glienke *et al*., 2022; Harrer *et al*., 2022; Fischer-Riepe *et al*., 2024; Umland *et al*., 2025). TRUCKs differ from synNotch and SNIPR in how antigen recognition is translated into transgene expression. In TRUCK systems, a CAR is coupled to a synthetic inducible expression cassette regulated by tandem NFAT response elements upstream of a minimal promoter (Sup. Fig. 1a) (Chmielewski and Abken, 2020). Upon antigen engagement, the CAR activates endogenous T-cell signaling pathways that culminate in NFAT-dependent transcription of the downstream synthetic transgene encoding the payload. Because of this reliance on endogenous T-cell signaling, TRUCKs are considered non-orthogonal systems.

**Figure 1.**
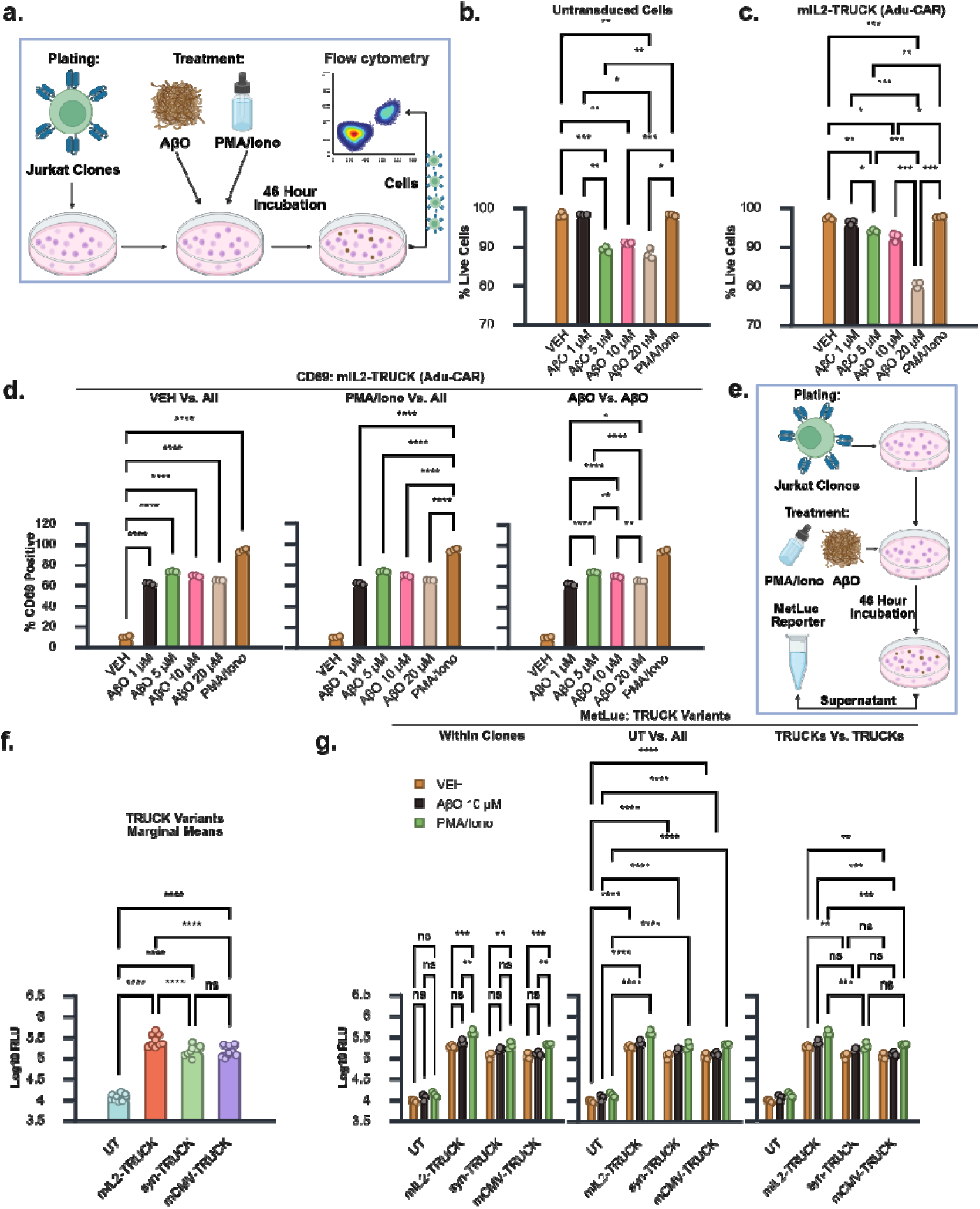
Human Aβ-targeting TRUCK systems contain functional receptors and inducible expression cassettes. **a.** Experimental design for flow cytometric analysis. After 46 h of AβO treatment, cells were harvested for assessment of viability and CD69 expression. **b**, **c**. Percentage of live cells following AβO treatment in (**b**) untransduced (UT) cells and (**c**) mIL2-TRUCK clones. Only statistically significant p-values are shown. **d.** Percentage of CD69-positive (CD69⁺) live mIL2-TRUCK cells. “VEH vs. All,” “PMA/Iono vs. All,” and “AβO vs. AβO” indicate specific comparisons derived from the same analysis for clarity of visualization. **e**. Experimental design for MetLuc analysis. Following 46 h of AβO treatment, conditioned media were collected for quantification of secreted MetLuc. **f**. Log10-transformed MetLuc RLU marginal means comparing UT, mIL2-, syn-, and mCMV-TRUCK clones. **g**. Log10 MetLuc RLU comparisons among UT and TRUCK variants at the indicated AβO concentrations. “Within Clones,” “UT vs. All,” and “TRUCKs vs. TRUCKs” indicate specific comparisons derived from the same analysis for clarity of visualization. Statistical analyses: (**b**, **c**) Games-Howell, (**d**) Tukey, and (**f**) Bonferroni all pairwise comparisons and (**g**) Bonferroni corrected planned comparisons. Full statistical analysis available in Table S2a-d. * p□<□0.05, ** p□<□0.01, *** p□<□0.001, **** p□<□0.0001. Dat are presented as mean□±□SEM from three biological replicates.

By contrast, synNotch and SNIPR are considered orthogonal systems because payload expression does not rely on endogenous signaling (Zhu *et al*., 2022; Morsut *et al*., 2016; Roybal *et al*., 2016). Instead, ligand engagement triggers regulated intramembrane proteolysis (RIP) that cleaves and releases an artificial transcription factor (ATF) present in the intracellular domain of the receptor (Sup. Fig. 1b) (Manhas *et al*., 2022). The ATF then translocates to the nucleus, where it binds its cognate synthetic promoter to drive transgene expression. Out of the two orthogonal receptor systems, synNotch was the first to be developed (Morsut *et al*., 2016; Roybal *et al*., 2016). However, it is not sensitive to monomeric soluble ligands and relies on large non-human RIP domains derived from mouse Notch1. Albeit less extensively studied, SNIPR was subsequently developed to overcome these limitations, resulting in a humanized, compact, and soluble ligand sensing platform (Zhu *et al*., 2022; Piraner *et al*., 2025).

In the context of AD, we recently showed that an Aβ-targeting synNotch receptor based on Aducanumab can detect extracellular Aβ aggregates and induce secretion of chimeric human–mouse versions of FDA-approved Aβ-clearing antibodies chLecanemab and Aducanumab (chAducanumab) *in vitro* (Bergo *et al*., 2025). In a recent preprint, others have generated Aβ-targeting synNotch receptors based on the Aβ-clearing antibody bapineuzumab, which was expressed in astrocytes to deliver brain-derived neurotrophic factor (BDNF), soluble TNF receptor (sTNFR1), and IL-1 receptor antagonist (IL-1Ra) *in vitro* (Spetz *et al*., 2025). The preprint also describes what to our knowledge is the first Aβ-targeting SNIPR. Albeit not in AD, *in vivo* proof of concept comes from a study using engineered CD4⁺ T cells with a synNotch that detected the brain extracellular matrix protein brevican (BCAN) and induced interleukin-10 (IL-10) production in response, reducing pathology in mouse models of multiple sclerosis (Simic *et al*., 2024).

Collectively, the literature establishes that synthetic receptors can feasibly support disease-responsive production of therapeutic payloads across diverse biological contexts. However, there are no direct comparisons of Aβ-targeting TRUCK, synNotch, and SNIPR systems. To address this gap, we performed a controlled head-to-head comparison of the receptor systems in human Jurkat cells exposed to oligomer-enriched Aβ (AβO) preparations. To isolate the contribution of receptor architecture itself, all three platforms incorporated the same Aducanumab-derived antigen-recognition domain, while synNotch and SNIPR additionally shared the ATF (Gal4VP64) and inducible expression cassette (Gal4UAS-mCMV). To compare the effects of different regulatory regions within the TRUCK system, we created three NFAT-responsive variants differing only in their minimal promoters (6xNFAT-mIL2, 6xNFAT-synTATA, and 6xNFAT-mCMV). Using Metridia Luciferase (MetLuc) as a shared inducible reporter in TRUCK, synNotch, and SNIPR, we found that extracellular AβO robustly induced output from synNotch and SNIPR, whereas none of the TRUCK architectures produced measurable MetLuc output above baseline. Among the orthogonal systems, SNIPR-driven MetLuc production outperformed synNotch, a finding confirmed by additional tests assessing chLecanemab production.

## Results

### An Aducanumab-derived scFv confers Aβ aggregate recognition to a human CAR

For the initial evaluation of TRUCK function, we generated a single lentiviral vector encoding both the inducible expression cassette and an Aβ-targeting CAR (Sup. Fig. 1c). The inducible cassette regulatory region consisted of a 6xNFAT-mIL2 (Table S1a), an architecture previously developed for NFAT-dependent reporter and therapeutic systems (Hooijberg *et al*., 2000; Chmielewski *et al*., 2011; Zhang *et al*., 2011). Specifically, the 6xNFAT sequence was derived from Sequence 7 of patent US 8556882 (GenBank: HJ489689.1) and the minimal IL-2 promoter sequence was obtained from pGL3-NFAT luciferase plasmid (RRID:Addgene_17870) (Clipstone and Crabtree, 1992). This regulatory module controlled a payload analogous to that employed in our synNotch studies in NIH-3T3 cells (Bergo *et al*., 2025), consisting of a single-chain scFv-Fc chLecanemab (Table S1b) and a secreted MetLuc separated by a T2A sequence (Sup. Fig. 1c).

For the CAR, we chose a second-generation architecture with a single co-stimulatory domain. The antigen-binding domain was adapted from an Aducanumab-derived single-chain variable fragment (scFv), previously validated by our group in CAR and synNotch configurations for the detection of Aβ aggregates in mouse DO11.10 and NIH-3T3 cells (Bergo *et al*., 2025; Siebrand *et al*., 2025). The scFv consists of the variable heavy (VH) and variable light (VL) chains of Aducanumab joined by a (G₄S)₃ flexible linker (Sup. Fig. 1c; Table S1c). The CAR backbone was selected to minimize ligand-independent and off-target activation and thereby reduce potential basal NFAT-driven payload expression. Although CD28 hinge and transmembrane regions (CD28-HT) can confer stronger signaling (Alabanza *et al*., 2017; Majzner *et al*., 2020), they have been associated with increased off-target activation relative to the same CARs using CD8α-based hinge and transmembrane regions (CD8-HT) (Muller *et al*., 2021). Consequently, the human Adu-CAR incorporated a CD8-HT, previously implemented in FMC63-based CD19 CARs and in the FDA-approved tisagenlecleucel (Sup. Fig. 1c; Table S1c) (Majzner *et al*., 2020; Muller *et al*., 2021).

Tisagenlecleucel employs a 4-1BB co-stimulatory domain; however, the present receptors are intended for application in Tregs rather than cytotoxic CD4⁺/CD8⁺ T cells. In human CAR-Tregs, CD28 co-stimulatory domains have been reported to better preserve lineage stability and functional activity than 4-1BB-based designs (Boroughs *et al*., 2019; Dawson *et al*., 2020; Lamarthée *et al*., 2021). For this reason, the CD8α hinge–transmembrane region was fused to a CD28 co-stimulatory domain, followed by a human CD3ζ signaling domain (Sup. Fig. 1c; Table S1c).

The initial tests focused on the validation of the Adu-CAR within the mIL2-6NFAT-based TRUCK system (mIL2-TRUCK). To do this, we resorted to using Jurkat cells, a widely used human CD4⁺ T-cell leukemia–derived cell line that has been previously used to validate NFAT-driven mIL2 reporter systems as well as CAR and TRUCK systems (Clipstone and Crabtree, 1992; Hooijberg *et al*., 2000; Alonso-Camino *et al*., 2013; Lipowska-Bhalla *et al*., 2013; Uchibori *et al*., 2019; Bloemberg *et al*., 2020).

*In vitro* models of human AD commonly rely on exogenous administration of Aβ to human primary neurons, a strategy widely used in the field (Busciglio, Lorenzo and Yankner, 1992; Mattson *et al*., 1992; Busciglio, Yeh and Yankner, 1993; Busciglio *et al*., 1995; Zhang *et al*., 2002, 2003; Jana and Pahan, 2010; Sivananthan *et al*., 2010; Jhamandas *et al*., 2011) adapted by our group (Bergo *et al*., 2025; Siebrand *et al*., 2025; Walton *et al*., 2025). We adopted this approach to evaluate activation of Adu-CAR-expressing Jurkat cells. Aβ monomers aggregate into toxic species found in AD senile plaques once they are released into the extracellular space after the proteolytic cleavage of the APP (Hampel *et al*., 2021). Thus, unlike most CAR targets in cancer, Adu-CAR’s target does not form at cell membranes but in the extracellular space as an aggregate. Accordingly, for the experiments, untransduced cells (UT) and cells expressing the complete mIL2-TRUCK construct in culture were treated with vehicle (VEH), oligomer-enriched Aβ (AβO; 1, 5, 10, or 20 µM), or phorbol 12-myristate 13-acetate plus ionomycin (PMA/Iono; Fig. 1a), used as a positive control for NFAT activation.

We first evaluated cell viability by analyzing the percentage of live cells across treatment conditions by flow cytometry (Sup. Fig. 1d). A Welch’s ANOVA revealed significant differences between treatments for untransduced cells (W(5, 5.14) = 256.400, p < 0.0001; Table S2a). Multiple comparisons showed that AβO concentrations of 5–20 µM resulted in significantly reduced viability relative to VEH (p ≤ 0.006) and PMA/Ionomycin (p ≤ 0.017; Fig. 1b). For mIL2-TRUCK–expressing cells there were also significant differences between groups (W(5, 5.19) = 275.080, p < 0.0001; Table S2b). Multiple comparisons demonstrated that AβO 5–20 µM treatments significantly reduced viability relative to VEH (p ≤ 0.048) and also relative to PMA/Ionomycin (p < 0.05; Fig. 1c). These findings are consistent with previously reported AβO-associated cytotoxicity (Cline *et al*., 2018) and align with our prior observations in DO11.10 cells with or without CAR expression (Siebrand *et al*., 2025).

CD69 expression was used to assess functional activation of the human Adu-CAR, as previously done by our group in DO11.10 cells (Siebrand *et al*., 2025) and by others in Jurkat models to validate CAR (Alonso-Camino *et al*., 2013; Lipowska-Bhalla *et al*., 2013; Bloemberg *et al*., 2020). Density plots confirmed detection of the HA epitope used to detect Adu-CAR in transduced cells under VEH and PMA/ionomycin conditions (Sup. Fig. 1e). However, HA signal was reduced in AβO-treated samples, consistent with receptor internalization, masking of the HA antibody epitope by Aβ aggregates, or both. Similar findings have been reported by others testing Aβ-targeting CARs in mouse CD4 T cells (Boskovic *et al*., 2026). Accordingly, subsequent analyses were performed on the total cell population (HA-positive and negative) within each treatment condition.

With the exception of PMA/Iono positive control treatment, the percentage of CD69-positive cells following AβO treatment was markedly higher in Adu-CAR–expressing Jurkat cells than in UT controls (Sup. Fig. 1f, g), focusing subsequent analyses on Adu-CAR–expressing cells. There were highly significant treatment-dependent differences in CD69 expression (F(5, 12) = 2560.202, p < 0.0001, η² = 0.9991; Table S2c). Post hoc comparisons showed that all AβO concentrations induced significantly greater CD69 expression relative to VEH (all p < 0.0001; Fig. 1d, VEH Vs. All). However, CD69 induction by AβO remained significantly lower than that observed with PMA/Ionomycin (all p < 0.0001; Fig. 1d, PMA/Iono Vs. All). Among AβO concentrations, the maximal response occurred at 5 µM (all p ≤ 0.0025; Fig. 1d, AβO Vs. AβO), with reduced activation at higher concentrations, consistent with toxicity-associated attenuation of signaling.

Having validated functional signaling of the human Adu-CAR, we generated two additional TRUCK variants to characterize the regulation of an inducible transgene by this receptor system. This was done to overcome potential limitations of the use of an mIL2 promoter in cells that actively suppress IL-2 production, such as Tregs. The variants were obtained by replacing the mIL2 promoter with either a synthetic TATA box promoter (synTATA; Table S1d), described in previous studies (Merlet *et al*., 2013; Zimmermann *et al*., 2020), or a minimal CMV promoter (mCMV; Table S1e), commonly used in synNotch and SNIPR systems (Morsut *et al*., 2016; Roybal *et al*., 2016; Zhu *et al*., 2022; Bergo *et al*., 2025). The resulting 6xNFAT-mIL2 (mIL2-TRUCK), 6xNFAT-synTATA (syn-TRUCK), and 6xNFAT-mCMV (mCMV-TRUCK) constructs all incorporated the same Adu-CAR receptor (Sup. Fig. 1c) and identical payload cassette (Sup. Fig. 1h).

The inducible expression cassette encodes the Aβ-targeting antibody chLecanemab (Sup. Fig. 1h). However, direct quantification of chLecanemab does not allow valid comparisons across all treatment conditions. Under VEH conditions, where antibody production is ligand-independent, all antibody produced by the cells remains soluble in the culture medium. In contrast, in the presence of AβO, the antibody partitions between soluble and aggregate-bound fractions in proportions that cannot be directly determined. As a result, ELISA-based measurement of soluble antibody under AβO conditions underestimates total production relative to VEH. Conversely, quantification of aggregate-bound antibody by immunocytochemistry cannot be compared to VEH conditions because, in the absence of Aβ, no antibody is retained on aggregates. Therefore, neither soluble nor aggregate-associated measurements alone accurately reflect total antibody production across treatments. This, in turn, prevents the direct comparison of receptor ligand-independent activation under VEH treatments relative to receptor ligand-dependent activation under AβO treatments.

For this reason, we used secreted MetLuc, co-expressed with chLecanemab from the same inducible cassette via a T2A sequence (Sup. Fig. 1h), as a soluble quantitative surrogate for total payload production. Unlike chLecanemab, MetLuc remains entirely soluble under all treatment conditions, enabling direct and unbiased comparison of receptor-driven output between VEH and AβO treatments. Thus, for these experiments, untransduced Jurkat controls and cells expressing mIL2-, syn-, or mCMV-TRUCK were treated for 46 h with VEH, oligomer-enriched AβO (10 µM), or PMA/ionomycin activity (Fig. 1e). Conditioned media were then collected and analyzed for MetLuc.

Following Log10 transformation of the raw MetLuc RLU values to meet the assumptions of normality (p ≥ 0.057) and homogeneity of variances (p = 0.350), we performed a two-way ANOVA to perform a joint analysis of mIL2-, syn-, and mCMV-TRUCK systems. The interaction between clone and treatment was not significant (F(6, 24) = 1.980, p = 0.109, partial η² = 0.331), whereas the main effects of clone (F(3, 24) = 947.002, p < 0.0001, partial η² = 0.992) and treatment (F(2, 24) = 54.876, p < 0.0001, partial η² = 0.821) were highly significant (Table S2d).

Because the interaction was not significant, marginal means were examined first. Marginal means represent the effect of one factor after averaging across the levels of the other factor. Bonferroni-corrected marginal means comparisons showed that UT cells produced less MetLuc than all TRUCK-expressing groups (all p < 0.0001; Fig. 1f). Among TRUCK variants, mIL2-TRUCK yielded higher MetLuc output than syn-TRUCK (p < 0.0001) and mCMV-TRUCK (p < 0.0001), whereas syn-TRUCK and mCMV-TRUCK did not differ (p = 1). Across treatments, VEH was associated with lower MetLuc than both AβO (p = 0.002) and PMA/ionomycin (p < 0.0001), and PMA/ionomycin produced higher output than AβO (p < 0.0001, Sup. Fig. 1i).

The clone marginal means described above are obtained by averaging MetLuc production across all treatment conditions within each clone; consequently, VEH RLU values contributed to these averages and lowered the overall estimates. Conversely, treatment marginal means were calculated by averaging across all clones for each condition, which incorporated RLU values from UT control cells and thereby reduced the apparent treatment effect. Because these averaged estimates do not isolate the biologically relevant comparisons, Bonferroni-corrected planned contrasts were examined despite the absence of a significant interaction. These contrasts were restricted to (i) comparisons between VEH, AβO and PMA/Iono within the same clone and (ii) comparisons between clones within the same treatment condition. This strategy avoided unnecessary correction for the familywise error rate for comparisons across different treatments and clones that were not experimentally informative.

As expected, UT cells showed no treatment-dependent differences (p ≥ 0.117; Fig. 1g, Within Clones). In contrast, PMA/Iono significantly increased MetLuc production relative to VEH for all TRUCK variants (p ≤ 0.001). Notably, AβO 10 µM did not produce a significant increase relative to VEH for any TRUCK system (p ≥ 0.106). Nevertheless, all TRUCK conditions, including VEH, remained significantly higher than UT controls (all p < 0.0001; Fig. 1g, UT vs. All), evidencing basal activation. Direct comparisons among TRUCK variants showed that syn-TRUCK and mCMV-TRUCK did not differ under any condition (p = 1; Fig. 1g, TRUCKs Vs. TRUCKs). Except for the AβO 10 µM condition (p = 0.069), mIL2-TRUCK produced higher MetLuc output than the other TRUCK systems across treatments (p ≤ 0.002).

Taken together, these findings indicate limited performance of TRUCK platforms as payload delivery systems under the conditions tested. This limited output cannot be attributed to dysfunctional Adu-CAR signaling, as Adu-CAR–expressing Jurkat cells demonstrate robust activation in response to Aβ treatment (Fig. 1d). Nor can it be explained by inactive regulatory elements, as all three 6xNFAT-based expression cassettes drive significantly increased MetLuc production following PMA/Iono stimulation (Fig. 1g), confirming their responsiveness to NFAT activation.

Nevertheless, albeit non-significant, there was a pattern of MetLuc production in response to 10 µM AβO across the different replicates for mIL2 and syn-TRUCK variants that was consistent with a potentially biologically relevant weak activation. Because of this, for the next studies we included the three TRUCK receptor variants in a direct comparison with Aducanumab-based SNIPR and synNotch systems.

### Aducanumab-based SNIPR and synNotch systems show qualitatively superior responses to Aβ aggregates compared to TRUCK systems

We have previously validated an Aβ-targeting Adu-synNotch receptor using the same Aducanumab scFv used for the Adu-CAR in the TRUCK systems tested above (Bergo *et al*., 2025). The regulatory core of synNotch receptors is derived from the mouse Notch1 transmembrane domain (Morsut *et al*., 2016; Roybal *et al*., 2016), whose proteolytic cleavage mechanism is highly conserved across species. As a result, the amino acid sequence of the Aducanumab scFv and the regulatory region of the synNotch receptor tested here in human cells are identical to that used in our prior studies in mouse cells (Sup. Fig. 2a; Table S1f). However, for consistency across receptor formats, the commonly used N-terminal Myc-tag was replaced with the same HA epitope employed in the Adu-CAR of the TRUCK systems tested above.

**Figure 2.**
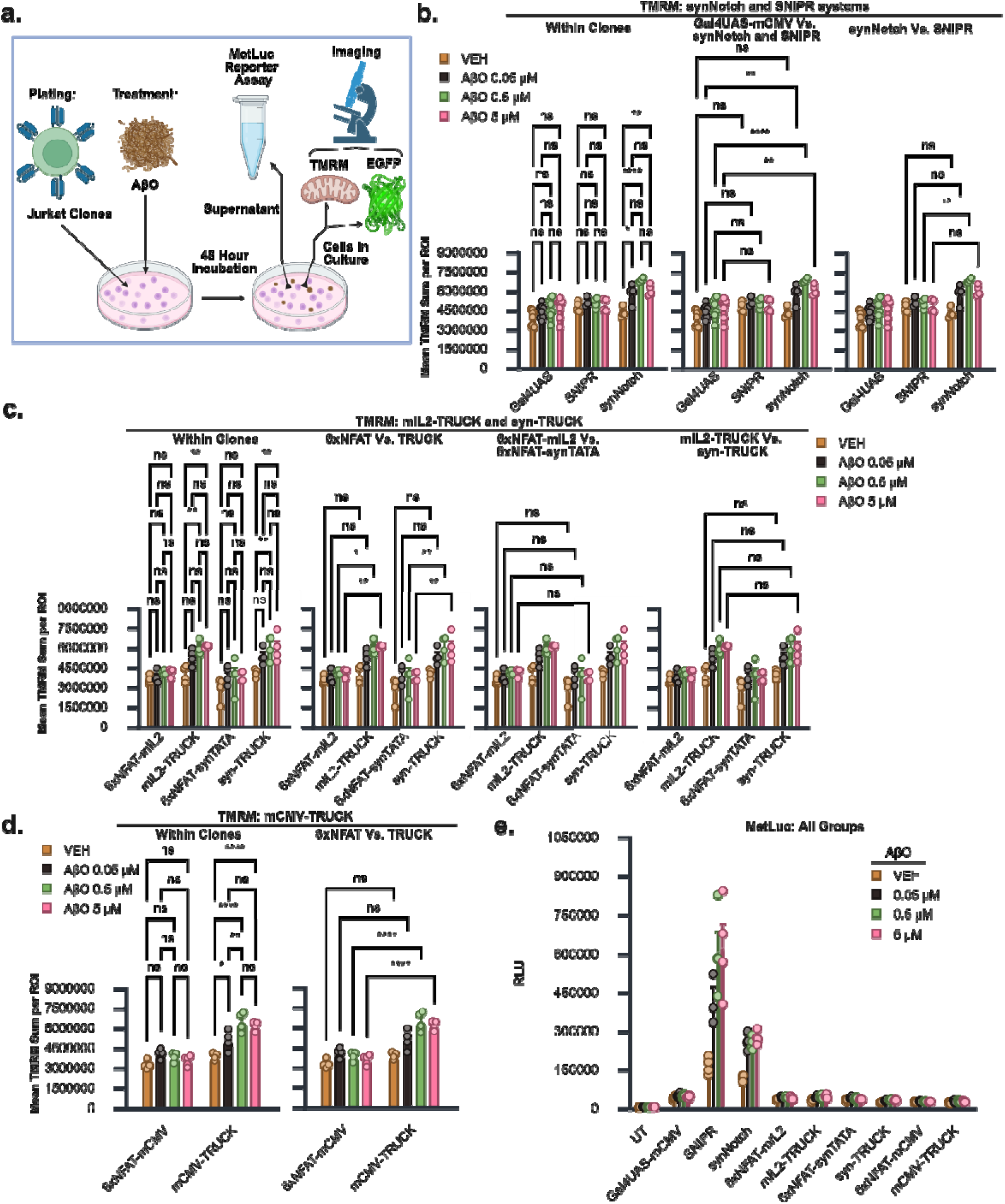
synNotch and SNIPR systems exhibit higher MetLuc output than TRUCK constructs in response to AβO. **a.** Experimental design for TMRM, EGFP, and MetLuc quantification following 48 h treatment with AβO. **b.** Mean TMRM sum comparisons assessing differences between synNotch and SNIPR systems at the indicated AβO concentrations. This metric serves as a proxy for the average total TMRM fluorescence per cell. “Within Clones,” “Gal4UAS-mCMV vs. synNotch and SNIPR,” and “synNotch vs. SNIPR” indicate specific comparisons derived from the same analysis for clarity of visualization. **c.** Mean TMRM sum comparisons assessing differences between mIL2-TRUCK and syn-TRUCK systems. “Within Clones,” “6xNFAT vs. TRUCK,” “6xNFAT-mIL2 Vs. 6xNFAT-synTATA”, and “mIL2-TRUCK vs. syn-TRUCK” indicate specific comparisons derived from the same analysis. **d.** Mean TMRM sum comparisons assessing the mCMV-TRUCK system. “Within Clones” and “6xNFAT vs. TRUCK” indicate specific comparisons derived from the same analysis. **e**. Raw MetLuc RLU shown for all groups. Data are presented a mean ± SEM from at least three independent biological replicates. Statistical analyses: (**b**, **c, d**) Bonferroni corrected planned comparisons. Full statistical analysis available in Table S3a-c. * p□<□0.05, ** p□<□0.01, *** p□<□0.001, **** p□<□0.0001. Data are presented as mean□±□SEM from at least three biological replicates.

For the SNIPR construct, we again used the same human CD8α signal peptide, HA-Tag, and Aducanumab scFv as in Adu-CAR and Adu-synNotch constructs (Sup. Fig. 2b; Table S1g). For the hinge and transmembrane domains, we adopted a receptor configuration incorporating human-derived Notch regulatory domains previously demonstrated to efficiently respond to soluble ligands (Piraner *et al*., 2025).

To enable a direct comparison between the receptor element of each system, the SNIPR and synNotch systems shared both the ATF and the inducible expression cassette sequence (Sup. Fig. 2a, b; Table S1f, g). In both platforms, the ATF consisted of the Gal4 DNA-binding domain fused to the VP64 transcriptional activator (Gal4VP64), which contains four tandem VP16 activation domains (Beerli *et al*., 1998). SNIPR and synNotch also employed an identical Gal4UAS-mCMV inducible regulatory architecture (Sup. Fig. 2a, b; Table S1h). Thus, the antigen-binding domain, ATF, and inducible expression cassettes were identical for both orthogonal receptor systems. Notably, the mCMV promoter sequence within this cassette was identical to that used in the 6xNFAT–mCMV TRUCK variant to improve the comparison of promoter behavior across orthogonal and non-orthogonal systems.

For these experiments, we analyzed clones expressing the complete TRUCK, SNIPR, and synNotch receptor systems alongside clones expressing only the corresponding inducible expression cassettes. To enable this comparison, synNotch (Sup. Fig. 2a), SNIPR (Sup. Fig. 2b), and TRUCK (Sup. Fig. 2c) constructs were separated into two independent lentiviral vectors, one for the receptor and another one for the inducible expression cassette. Jurkat cells transduced with receptor-encoding vectors were identified by HA-tag expression, whereas cells transduced with inducible expression cassette vectors were identified by enhanced green fluorescent protein (EGFP), driven by a separate constitutive promoter within the same vector. This configuration enabled fluorescence-activated cell sorting (FACS) using consistent gating parameters to sort receptors (HA-tag) and expression cassettes (EGFP) within standardized levels of expression to minimize variability owed to transduction efficiencies (Sup. Fig. 2d, e).

Because maximal Adu-CAR activation occurred at 5 µM AβO (Fig. 1d), the cells in this new MetLuc experiment were treated for 48 hours with oligomer-enriched AβO at 0.05 µM, 0.5 µM, or 5 µM, or with VEH control, after which conditioned media were collected and analyzed (Fig. 2a). This design enabled dissection of distinct signaling components. Basal transcriptional leakage (TL) was quantified using clones expressing the inducible expression cassette alone. Receptor ligand-independent activation (LIA) was assessed in full receptor systems under VEH treatment (LIA + TL), and receptor ligand-dependent activation (LDA) was determined by comparing AβO-treated conditions (LDA + LIA + TL) to their corresponding VEH controls (LIA + TL).

Additionally, we evaluated Tetramethylrhodamine, methyl ester (TMRM) and EGFP signal (Fig. 2a) in live cultured cells prior to media collection utilizing fluorescent imaging (Sup. Fig. 2f). TMRM is a cell-permeant dye that accumulates in viable cells with active mitochondria (Ehrenberg *et al*., 1988; Farkas *et al*., 1989), and mitochondrial membrane potential increases with T cell activation, with higher ΔΨm associating with greater effector activity including Granzyme B expression and proliferation (Sukumar *et al*., 2016; Amitrano *et al*., 2021), allowing us to simultaneously gain insights into viability and relative activation to contextualize synNotch, SNIPR, and TRUCK receptor activation.

The mean sum of TMRM per cell TMRM signal intensity was relatively homogeneous across UT and expression cassette-only clone (Sup. Fig. 2g). With the exception of SNIPR clones, all receptor-bearing clones displayed a clear pattern of increased signal relative to UT. To formally assess differences in TMRM, we separately analyzed the orthogonal systems (synNotch and SNIPR) and the non-orthogonal systems (TRUCKs).

We identified an extreme outlier in one replicate of the SNIPR AβO 0.05 µM treatment group, which was 3.01 interquartile ranges below Quartile 1 of the SNIPR clones. The reason for this group’s reduced viability, as indicated by TMRM values, was unclear; therefore, it was excluded from all subsequent analyses to ensure that the remaining groups accurately represented the viability and health of that particular clone across treatments for receptor system comparisons. Subsequent analysis revealed a significant interaction between clone and AβO treatment (F(6, 51) = 3.083, p = 0.0117, partial η² = 0.2662; Table S3a). Planned comparisons showed no significant differences between treatments within clones expressing only the Gal4UAS-mCMV inducible cassette or the SNIPR clones (p ≥ 0.3388; Fig. 2b, Within Clones), nor when comparing said clones across the same treatments (p ≥ 0.5805, Fig. 2b, Gal4UAS-mCMV vs. synNotch and SNIPR). However, synNotch-expressing clones exhibited a significant increase in TMRM signal between 0.05, 0.5 and 5 µM AβO treatments relative to VEH clones (p ≤ 0.0208; Fig. 2b, Within Clones), as well as a significant difference when comparing AβO 0.05 to 5 µM treatments against Gal4UAS-mCMV clones (p ≤ 0.0083), but not with VEH (p = 1, Fig. 2b, Gal4UAS-mCMV vs. synNotch and SNIPR). Despite this, there were only significant differences when directly comparing the same AβO 0.5 µM treatments between synNotch and SNIPR (p = 0.0026; Fig. 2b, synNotch Vs. SNIPR).

The combined analysis of the mIL2, syn, and mCMV-TRUCK systems revealed a violation of the homogeneity of variances (Levene’s test, p = 0.028), which could not be remedied through data transformation. Subsequently, we focused on a joint analysis comparing the two TRUCK systems previously validated in the literature, mIL2 and syn-TRUCKs (Chmielewski *et al*., 2011; Chmielewski and Abken, 2017, 2020; Kunert *et al*., 2018; Liu *et al*., 2019; Zimmermann *et al*., 2020; Glienke *et al*., 2022; Harrer *et al*., 2022; Fischer-Riepe *et al*., 2024; Umland *et al*., 2025), which met the assumption of homogeneity of variances (Levene’s test, p = 0.065).

A two-way ANOVA did not show a significant interaction (F(9, 48) = 1.242, p = 0.2929, partial η² = 0.1889), but significant main effects were observed for clone (F(3, 48) = 31.010, p < 0.0001, partial η² = 0.6596) and AβO treatment (F(3, 48) = 14.642, p < 0.0001, partial η² = 0.4778; Table S3b). Given the absence of an interaction, we concentrated on the analysis of the main effects first. Multiple comparisons of the clone marginal means indicated that the 6xNFAT-mIL2 and 6xNFAT-synTATA clones were not significantly different from each other (p = 1; Sup. Fig. 2h), while both expression cassette-only clones exhibited significantly lower activity compared to their mIL2-TRUCK (p < 0.0001) and syn-TRUCK (p < 0.0001) counterparts (Sup. Fig. 2h). However, there were no statistically significant differences between the mIL2-TRUCK and syn-TRUCK clones (p = 1; Sup. Fig. 2h). Regarding the main effects of AβO, the marginal means for VEH were significantly lower than those for the other treatments (p ≤ 0.001), but no differences in TMRM levels were observed between the AβO 0.05-5 µM treatments (p ≥ 0.414; Sup. Fig. 2i).

We followed up with Bonferroni-corrected planned comparisons to explore potentially biologically relevant patterns (Fig. 2c). Both mIL-2 and syn-TRUCKs presented significantly higher TMRM at AβO 0.5 and 5 µM when compared to VEH (p ≤ 0.0093) but there were no differences between the 6xNFAT-mIL2 and 6xNFAT-synTATA groups (all p = 1; Fig. 2c, Within Clones), an activation pattern that parallels that of synNotch and Gal4UAS-mCMV (Fig. 2b, Within Clones). Relative to their 6xNFAT-expressing only controls, both mIL2-TRUCKS (p ≤ 0.0149) and syn-TRUCKs (p ≤ 0.0016) showed significantly higher TMRM only at AβO 0.5 and 5 µM (all p ≤ 0.0149; Fig. 2c, 6xNFAT Vs. TRUCK). There were no significant differences when comparing 6xNFAT-mIL2 and 6xNFAT-synTATA-only clones (all p = 1; Fig. 2c, 6xNFAT-mIL2 Vs. 6xNFAT-synTATA) nor when comparing the mIL2- and syn-TRUCK clones (all p = 1; Fig. 2c, mIL2-TRUCK Vs. syn-TRUCK).

The assumption of homogeneity of variances was satisfied for the remaining mCMV-TRUCK and 6xNFAT-mCMV groups (Levene’s test, p = 0.123). The omnibus test revealed a significant interaction between clone and AβO treatment (F(3, 24) = 9.312, p = 0.0003, partial η² = 0.5379; Table S3c). Planned comparisons showed that mCMV-TRUCKs exhibited significantly higher TMRM levels at AβO concentrations of 0.05, 0.5, and 5 µM compared to the VEH control (p ≤ 0.0303; Fig. 2d, Within Clones). In contrast, there were no significant differences between treatments within the 6xNFAT-mCMV clones (p ≥ 0.3749; Fig. 2d, Within Clones). Similar to mIL2- and syn-TRUCKs (Fig. 2c), mCMV-TRUCKs demonstrated significantly higher TMRM compared to the control 6xNFAT-mCMV groups at AβO concentrations of 0.5 and 5 µM (p < 0.0001; Fig. 2d, 6xNFAT vs. TRUCK).

In summary, no concentration-response increase in TMRM was observed for Gal4UAS-mCMV, 6xNFAT-mIL2, 6xNFAT-synTATA, or 6xNFAT-mCMV (Fig. 2b-d). However, a concentration-response increase in TMRM was noted for synNotch, mIL2-TRUCKs, syn-TRUCKs, and mCMV-TRUCKs (Fig. 2b-d), while this response was absent in SNIPR clones (Fig. 2b). This discrepancy may be linked to the activation mechanism of SNIPR. For soluble ligands, SNIPR is internalized along with its cognate ligand to trigger the release of the ATF (Piraner *et al*., 2025). In contrast, the activation of CARs in TRUCKs and synNotch receptors does not involve internalization of the receptor-ligand complex, although endocytosis is not ruled out.

For the analysis of EGFP in the orthogonal receptors, there was only a significant effect of clone (F(2, 51) = 4.125, p = 0.02186, partial η² = 0.1392; Table S3d). The comparisons between marginal means revealed that the only difference was lower total EGFP in Gal4UAS-mCMV wells relative to synNotch (p = 0.025, Sup. Fig. 2j). However, planned multiple comparisons between clone and treatment combinations did not reveal any significant differences (all p = 1, Sup. Fig. 2k).

All assumptions were met for the combined analysis of all TRUCK systems, with a two-way ANOVA again only producing a significant main effect for clones (F(5, 72) = 6.009, p = 0.0001, partial η² = 0.2944; Table S3e). There were no differences in EGFP marginal means when comparing 6xNFAT-mIL2 with mIL2-TRUCKS (p = 1) or 6xNFAT-mCMV with mCMV-TRUCKS (p = 0.250), but 6xNFAT-synTATA clones had significantly higher values than syn-TRUCKs (p = 0.002; Sup. Fig. 2l). However, there were no significant differences among mIL2-TRUCKS, syn-TRUCKs, and mCMV-TRUCKS (p = 1; Sup. Fig. 2l) and planned comparisons did not show any significant differences between any clone and treatment combination (all p ≥ 0.1219; Sup. Fig. 2m).

Taken together, there was a distinct pattern for increased TMRM relative to inducible expression cassette-only clones in response to AβO for all receptor systems except SNIPR. This may indicate reduced viability, reduced activation, or both, which could negatively impact receptor function in the comparisons that follow. For EGFP, while differences were found when averaging all treatments by clones in marginal means, there were no differences when comparing clone and treatment combinations, which could otherwise significantly bias the analysis of MetLuc between systems.

Comparison of MetLuc RLU values across all treatment conditions showed that SNIPR- and synNotch-based systems generated signals more than an order of magnitude higher than those observed for TRUCK platforms (Fig. 2e; Sup. Fig 2n). Within the TRUCK systems, MetLuc production from clones expressing the full receptor construct was comparable to that of clones expressing only the inducible expression cassette, indicating minimal separation between receptor-driven and cassette-driven output (Sup. Fig 2n, MetLuc: TRUCKs). In contrast, receptor-expressing SNIPR and synNotch clones displayed markedly increased MetLuc output relative to their corresponding expression cassette–only controls (Sup. Fig 2n, MetLuc: SNIPR & synNotch), consistent with receptor-dependent activation.

Given the large differences in MetLuc output between platforms, TRUCK and SNIPR/synNotch systems were analyzed separately to avoid inflating model complexity and imposing unnecessary multiple-comparison penalties on the familywise error rate.

### TRUCK variants were not responsive to Aβ treatments

Basal activity of the regulatory regions (transcriptional leakage; TL) in the TRUCK expression cassettes is mechanistically plausible, given their reliance on endogenous NFAT signaling, which may exhibit some degree of constitutive activity in Jurkat cells. In contrast, comparable TL from the Gal4UAS-mCMV regulatory architecture used in SNIPR and synNotch systems would not be predicted if the system were fully orthogonal to endogenous transcriptional networks.

To formally assess whether the 6xNFAT system is not as tightly regulated as the orthogonal Gal4UAS systems, VEH-treated expression cassette–only clones for TRUCK, synNotch, SNIPR, and UT cells were analyzed using a one-way ANOVA. This analysis revealed significant differences in MetLuc RLU across groups (F(5, 18) = 115.626, p < 0.001, η² = 0.9605; Table S4a). All-pairwise multiple comparisons showed that all expression cassette–only clones produced significantly higher MetLuc RLU than UT cells (all p < 0.0001; Fig. 3a). Among transduced clones, 6xNFAT–mCMV exhibited significantly lower basal MetLuc output than all other expression cassettes (p ≤ 0.0016). No significant difference was detected between 6xNFAT–mIL2 and 6xNFAT–synTATA constructs (p = 0.9962). Gal4UAS-mCMV exhibited significantly higher basal activity than 6xNFAT–mIL2 (p = 0.0051) and 6xNFAT–synTATA (p = 0.002). Collectively, these results demonstrate that all transcriptional regulatory constructs drive basal payload expression in the absence of receptors. Unexpectedly, the orthogonal Gal4UAS-mCMV architecture did not exhibit tighter basal regulation than the non-orthogonal 6xNFAT-based systems in Jurkat cells.

**Figure 3.**
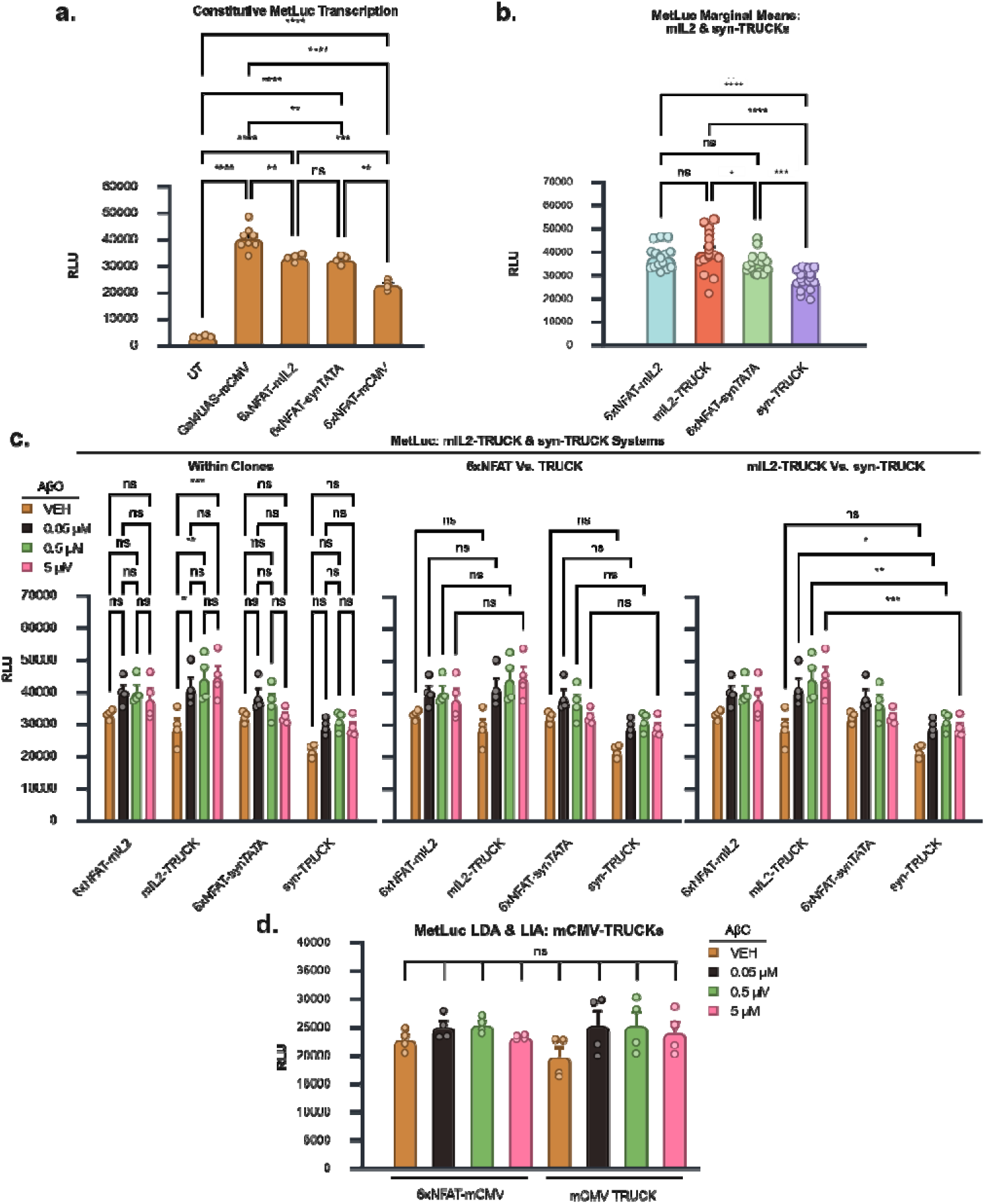
Human Aβ-targeting TRUCK systems do not produce MetLuc above basal transcriptional leakage in response to AβO. **a.** Comparison of basal transcriptional leakage (TL) under VEH conditions across orthogonal (Gal4UAS-mCMV) and non-orthogonal (6×NFAT) inducible expression cassettes relative to untransduced (UT) controls**. b.** Marginal means of MetLuc production for mIL2-TRUCK and syn-TRUCK full receptor systems compared with their corresponding 6×NFAT expression cassette–only controls. **c.** Planned comparisons of MetLuc RLU for mIL2- and syn-TRUCK systems. “Within Clones,” “6×NFAT vs. TRUCK,” and “mIL2-TRUCK vs. syn-TRUCK” denote specific contrasts derived from the same analysis for visualization purposes. **d.** All pairwise comparisons of MetLuc RLU within the mCMV-TRUCK system. The comparison bar denotes all possibl comparisons. Statistical analyses: (**a**) Tukey, (**b**) Bonferroni, and (**d**) Games-Howell all pairwise comparisons and (**c**) Bonferroni corrected planned comparisons. Full statistical analysis available in Table S4a-c. * p□<□0.05, ** p□<□0.01, *** p□<□0.001, **** p□<□0.0001. Data are presented as mean□±□SEM from four biological replicates.

Having characterized basal transcriptional leakage in the inducible expression cassettes, we next evaluated full TRUCK receptor systems to determine whether incorporation of the Adu-CAR conferred receptor-dependent activation beyond TL cassette-driven output. To formally assess differences in off-target receptor LIA and on-target receptor LDA between systems, expression cassette–only clones and full receptor system clones were analyzed in parallel. Under this framework, comparison of a full receptor system at a given AβO concentration (LDA + LIA + TL) with its corresponding VEH condition (LIA + TL) isolates receptor-mediated LDA ([LDA + LIA + TL] - [LIA + TL] = LDA). Comparison of the expression cassette–only clone (TL) with the corresponding full receptor system under VEH conditions (LIA + TL) assesses the contribution of LIA within that system ([LIA + TL] - [TL] = LIA). Finally, comparison of the expression cassette–only clone (TL) with the corresponding full receptor system at a given AβO concentration (LDA + LIA + TL) assesses the combined contribution of LIA and LDA within that system ([LDA + LIA + TL] - [TL] = LDA+LIA).

For analysis of the TRUCK systems using a single two-way ANOVA, assumption testing indicated a violation of homogeneity of variances for MetLuc RLU values (Levene’s test, p = 0.005) that could not be resolved by transformations. However, separating the more widely used mIL2-TRUCK and synTATA-TRUCK systems into one analysis and the mCMV-TRUCK system into a different analysis complied with ANOVA assumptions.

For the combined analysis of mIL2-TRUCK and synTATA-TRUCK systems, the interaction between clone and AβO treatment was not statistically significant (F(9, 48) = 1.509, p = 0.172, partial η² = 0.221; Table S4b). In contrast, both the main effect of clone (F(3, 48) = 21.132, p < 0.0001, partial η² = 0.569) and the main effect of AβO treatment (F(3, 48) = 13.123, p < 0.0001, partial η² = 0.451) were highly significant. Given the absence of a significant interaction and the presence of significant main effects, the marginal means were examined first.

Bonferroni-corrected marginal means comparisons revealed that synTATA-TRUCK clones produced significantly lower MetLuc RLU than all other clone conditions (all p ≤ 0.0003; Fig. 3b). In contrast, average MetLuc production by cells expressing the complete mIL2-TRUCK system did not differ significantly from that of clones expressing only the corresponding 6xNFAT–mIL2 expression cassette (p = 0.592). When averaging across clones, marginal means for AβO treatment indicated that VEH conditions produced significantly less MetLuc than all AβO concentrations tested (all p ≤ 0.0004), with no significant differences observed between AβO concentrations (all p ≥ 0.621; Sup. Fig. 3a).

Bonferroni-corrected comparisons revealed no significant differences between VEH and AβO treatments (0.05, 0.5, or 5 µM) for clones expressing 6xNFAT–mIL2 (all p = 1), 6xNFAT–synTATA (all p = 1), or synTATA-TRUCK (all p ≥ 0.298; Fig. 3c, Within Clones). Significant treatment-dependent increases were detected only for mIL2-TRUCK–expressing cells, which exhibited higher MetLuc RLU under AβO 0.05 µM (p = 0.017), 0.5 µM (p = 0.0011), and 5 µM (p = 0.0006) relative to VEH. However, direct comparisons between mIL2-TRUCK and 6xNFAT–mIL2 clones under matched treatment conditions did not reveal statistically significant differences (all p = 1; Fig. 3c, 6xNFAT Vs. TRUCK), indicating the lack of a functionally relevant response at the tested AβO concentrations. Surprisingly, the output by the syn-TRUCK system relative to its expression cassette only hinted at a suppression of MetLuc production by Adu-CAR, albeit none of the pairwise comparisons reached significance (p ≤ 0.0973; Fig. 3c, 6xNFAT Vs. TRUCK). Notably, although mIL2-TRUCK and synTATA-TRUCK clones did not differ under VEH conditions, mIL2-TRUCK clones produced significantly higher MetLuc RLU than synTATA-TRUCK clones at all AβO concentrations tested (all p ≤ 0.0233; Fig. 3c, mIL2-TRUCK Vs. syn-TRUCK).

For the remaining mCMV-TRUCK system, assumption testing for a two-way ANOVA indicated that normality was satisfied (p ≥ 0.066) but homogeneity of variances was violated (p < 0.0001). To accommodate this violation, groups were collapsed into a single factor and analyzed using a Welch’s one-way ANOVA, which does not assume equal variances. The omnibus test was not statistically significant (W(7, 9.41)= 1.660, p = 0.230; Table S4c). Consistent with this result, exploratory Games–Howell all-pairwise comparisons did not reveal any significant differences between groups (all p ≥ 0.285; Fig. 3d).

Overall, all inducible expression cassettes exhibited constitutive transcriptional activation of the MetLuc reporter (Fig. 3a). Unexpectedly, basal transcriptional leakage (TL) from the 6xNFAT-based regulatory elements was not greater than that observed for the orthogonal Gal4UAS- mCMV system (Fig. 3a). Out of all the TRUCK systems, only the mIL2-TRUCK system provided evidence of receptor LDA when comparing VEH to AβO treatments (Fig. 3c, Within Clones). However, despite the presence of a functionally validated Adu-CAR (Fig. 1d) and transcriptionally competent 6xNFAT regulatory sequences (Fig. 1g; Fig. 3a), none of the clones expressing the complete TRUCK system displayed MetLuc production that was significantly higher than treatment-matched expression cassette-only clones (Fig. 3c, 6xNFAT Vs. TRUCK; Fig. 3d).

Notably, although not statistically significant, VEH-treated full TRUCK clones consistently exhibited lower MetLuc output than their corresponding VEH-treated expression cassette–only controls across mIL2-, syn-, and mCMV-TRUCK systems (Fig. 3c, 6xNFAT Vs. TRUCK; Fig. 3d). To better visualize the receptor-associated contributions analyzed above, we subtracted MetLuc values from expression cassette–only clones (TL) from those of the matched full receptor clones under each treatment (Sup. Fig. 3b). This normalization revealed a negative receptor-associated output trend under VEH conditions across all TRUCK variants (i.e., negative LIA), consistent with reduced NFAT-driven transcription in the presence of the CAR. However, with increasing AβO concentrations, receptor-associated output progressively increased (Sup. Fig. 3b). Together, these observations may be consistent with a model in which CAR expression is associated with reduced basal NFAT activity under VEH conditions, and that antigen engagement partially restores or offsets this suppression, resulting in modest increases in apparent receptor-dependent output relative to cassette-only levels.

### Aducanumab-based SNIPR performs better than synNotch in response to A**β**O treatments

MetLuc RLU production for synNotch and SNIPR was visibly higher than the TRUCK systems (Fig. 2e), prompting the separate analysis of the orthogonal synthetic receptor systems. Although synNotch and SNIPR share the same Gal4UAS-mCMV inducible expression cassette, we initially maintained separate Gal4UAS expression cassette–only control groups (n = 4 each) for each receptor system. This design ensured that, if synNotch and SNIPR had required independent two-way ANOVAs due to violations of normality or homogeneity of variance assumptions, each receptor platform would retain its own matched control group without reusing data across analyses. However, after Log10 transformation, all assumptions for a joint two-way ANOVA were satisfied (Table S5a), permitting the two Gal4UAS expression cassette–only control groups to be combined into a single control group (n = 8) for the unified analysis. As noted above, one SNIPR replicate at 0.05 µM AβO was excluded because it was an extreme low outlier TMRM value, resulting in n = 3 for that condition.

A two-way ANOVA revealed a significant interaction between clone and AβO treatment on Log10 transformed MetLuc RLU (F(6, 51) = 20.606, p < 0.0001, partial η² = 0.7071; Table S5a). Planned pairwise comparisons revealed no significant differences among Gal4UAS-mCMV expression cassette–only clones across AβO treatment conditions (all p = 1; Fig. 4a, Within Clones), indicating the absence of ligand-dependent effects in the absence of a receptor. In contrast, SNIPR-expressing clones exhibited a significant increase in MetLuc production for all AβO concentrations (0.05, 0.5, and 5 µM) relative to VEH (all p < 0.0001), with no significant differences detected between AβO concentrations (all p ≥ 0.0564; Fig. 4a, Within Clones). Similarly, synNotch-expressing clones showed a robust increase in MetLuc output for all AβO treatments compared to VEH (all p < 0.0001), without differences among AβO concentrations (all p = 1; Fig. 4a, Within Clones). Direct comparisons between Gal4UAS-mCMV expression cassette–only clones with receptor-expressing SNIPR or synNotch clones under matched treatment conditions revealed highly significant differences in all cases (all p < 0.0001; Fig. 4a, Gal4UAS-mCMV Vs. synNotch and SNIPR), consistent with receptor-dependent activation. Comparisons between SNIPR and synNotch systems showed no significant difference under VEH conditions (p = 0.176), whereas SNIPR produced significantly higher MetLuc output than synNotch at all AβO concentrations tested (all p ≤ 0.0094; Fig. 4a, synNotch Vs. SNIPR).

**Figure 4.**
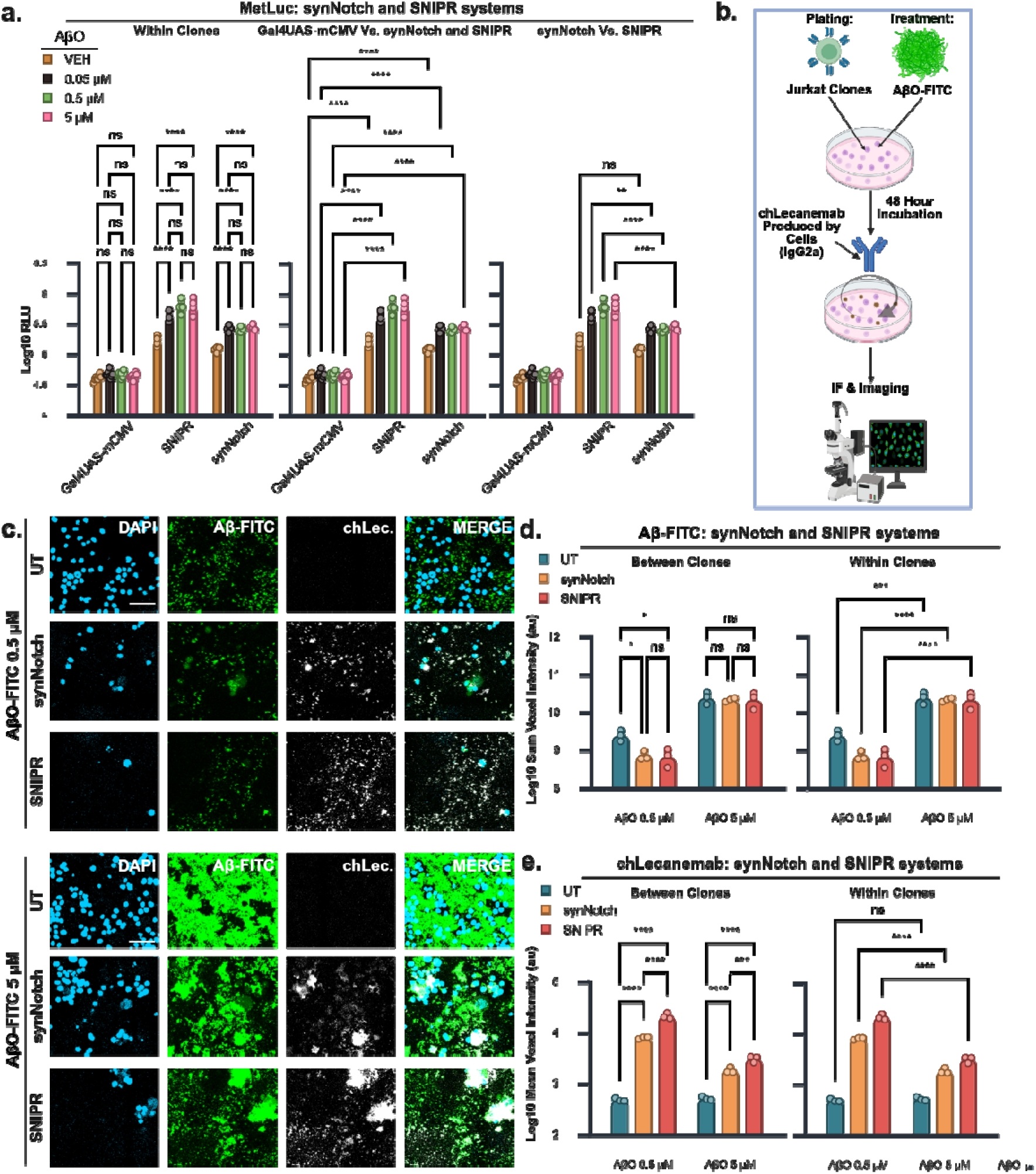
Human Aβ-targeting synNotch and SNIPR systems exhibit ligand-dependent activation, with SNIPR demonstrating superior output. **a.** Planned comparisons of Log10-transformed MetLuc RLU for synNotch and SNIPR systems. “Within Clones,” “Gal4UAS–mCMV vs. synNotch and SNIPR,” and “synNotch vs. SNIPR” denote specific contrasts derived from the same two-way ANOVA for visualization purposes. **b.** Experimental design for AβO–FITC treatments used to quantify Aβ and chLecanemab by immunofluorescence (IF) and confocal microscopy following 48 h exposure. **c.** Representative confocal images of untransduced (UT), synNotch-, and SNIPR-expressing clones under the indicated treatment conditions. Channels: DAPI (nuclei), Aβ–FITC (aggregates), and chLec. (chLecanemab). Images acquired using a 20× objective (1× zoom). Scale bar, 50 microns. Larger tile scans are shown in Sup. Fig. 4c. **d, e.** Planned comparisons of (**d**) summed Log10 Aβ–FITC signal and (**e**) mean chLecanemab voxel intensity within Aβ–FITC puncta for UT, synNotch-, and SNIPR-expressing clones. “Between Clones” and “Within Clones” indicate specific contrasts derived from the same statistical analysis for visualization clarity. Statistical analyses: (**a, d, e**) Bonferroni corrected planned comparisons. Full statistical analysis available in Table 5a-c. p□<□0.05, ** p□<□0.01, *** p□<□0.001, **** p□<□0.0001. Data are presented as mean□±□SEM from at least three biological replicates.

Based on the data presented above, both SNIPR and synNotch exhibited receptor LDA in response to AβO, with MetLuc production plateauing at 0.05 µM. SNIPR and synNotch presented comparable receptor LIA in the absence of AβO treatments. However, direct comparison between these two platforms revealed that SNIPR consistently produced higher ligand-dependent MetLuc output than synNotch at all AβO concentrations.

For the experiments above, MetLuc and chLecanemab were co-expressed from a 2A-based inducible bicistronic cassette, allowing MetLuc to serve as a quantitative surrogate for chLecanemab production and enabling direct comparisons of receptor output between VEH and AβO conditions. We next assessed chLecanemab production directly. To this end, a modified Gal4UAS-mCMV inducible expression cassette was generated in which the T2A–MetLuc module was removed, leaving chLecanemab as the sole transgene (Sup. Fig. 4a, b). The synNotch and SNIPR receptor constructs were identical to those used in the MetLuc experiments and were again delivered using a two–lentiviral vector configuration.

For these experiments, fluorescein-labeled Aβ(1–42) (Aβ–FITC) was used to enable the visualization of the target antigen directly. Aβ–FITC was incubated at 37 °C for 3 h to generate oligomer-enriched preparations (AβO–FITC), which were applied at final concentrations of 0.5 or 5 µM to untransduced, synNotch-, or SNIPR-transduced Jurkat cultures (Fig. 4b). After 48 h, cells were fixed and processed for immunofluorescence (IF) to detect chLecanemab colocalization with AβO–FITC.

The Fc domain of chLecanemab is derived from mouse IgG2a (Sup. Fig. 4a,b; Table S1b), enabling selective detection using an anti-mouse IgG secondary antibody by IF as previously described (Bergo *et al*., 2025). Cell nuclei were counterstained with DAPI. We observed a reduction in DAPI-positive nuclei for transduced cells relative to UT (Fig. 4c; Sup. Fig. 4c). This difference may reflect reduced cell recovery after washes during IF, decreased cell viability in the transduced populations, or both. Aβ–FITC signal appeared predominantly as discrete puncta at 0.5 µM and as larger aggregates at 5 µM (Fig. 4c; Sup. Fig. 4c). In synNotch- and SNIPR-transduced cultures, Aβ–FITC signal intensity appeared reduced relative to UT cells but, as expected, it was accompanied by clear chLecanemab signal.

For voxel-intensity analysis, 3D regions of interest (ROIs) were generated from Aβ–FITC puncta and used to quantify both Aβ–FITC and chLecanemab signals. Because EGFP and Aβ–FITC were detected in the same fluorescence channel, ROIs based on total signal would have incorporated EGFP-positive regions that do not colocalize with chLecanemab, artificially reducing the measured antibody signal. Therefore, although EGFP fluorescence was weak after PFA fixation, we minimized this potential bias by excluding nuclear and perinuclear EGFP-positive areas. Specifically, ROIs were restricted to Aβ–FITC puncta located ≥1 µm from DAPI-positive nuclei, thereby removing nuclear/perinuclear EGFP contribution from the analysis (Sup. Fig. 4d, e).

As a measure of total Aβ signal, a two-way ANOVA of summed Aβ-FITC intensity revealed a significant interaction between treatment and clone (F(2,12) = 4.392, p = 0.0371, partial η² = 0.4226; Table S5b). Notably, at 0.5 µM, Aβ-FITC signal was significantly lower in synNotch- and SNIPR-expressing cells compared to UT controls (p ≤ 0.0206; Fig. 4d, Between Clones). As expected, the sum Aβ-FITC intensity increased significantly from 0.5 to 5 µM AβO–FITC across all groups (p ≤ 0.0001; Fig. 4d, Within Clones).

To analyze antibody production across clones, we next assessed the mean chLecanemab signal intensity within Aβ-FITC puncta. A two-way ANOVA revealed a significant clone by treatment interaction (F(2,12) = 144.659, p < 0.0001, partial η² = 0.9602; Table S5c). At both concentrations, SNIPR-expressing cells exhibited higher chLecanemab signal than synNotch (p ≤ 0.0005) and UT cells (p < 0.0001), and synNotch exceeded UT (p < 0.0001; Fig. 4e, Between Clones). No treatment-dependent differences were observed in UT cells (p = 1, Fig. 4e, Within Clones). In contrast, both synNotch and SNIPR clones showed significantly lower mean chLecanemab intensity at 5 µM compared to 0.5 µM AβO–FITC (p < 0.0001, Fig. 4e, Within Clones). This reduction is consistent with the plateau observed in the MetLuc assays (Fig. 4a), indicating that, at higher antigen concentrations, a similar amount of antibody is distributed across a larger aggregate area, thereby reducing mean antibody signal intensity per aggregate.

Consistent with the MetLuc results, the IF analysis confirms receptor-dependent chLecanemab production for both synNotch and SNIPR systems as well as superior antigen-dependent output of SNIPR relative to synNotch.

## Discussion

This study shows that Aβ-targeting SNIPR and synNotch receptors outperform matched TRUCK architectures for inducible payload delivery in human Jurkat cells. Although the Adu-CAR used in the TRUCK constructs responded to AβO and all 3 6xNFAT inducible expression cassettes were functional, none of the TRUCK variants produced payload significantly above their matched expression cassette-only controls. By contrast, both orthogonal systems generated strong Aβ-dependent output with SNIPR consistently exceeding synNotch. The same pattern was seen in the MetLuc assays and in semi-quantitative antibody readout by IF, where SNIPR produced higher chLecanemab signal than synNotch.

### Basal transcriptional leakage

Off-target activation of synthetic receptors such as TRUCKs, synNotch, and SNIPRs is commonly assessed by comparing vehicle-treated and ligand-treated cells, an approach that captures basal receptor activation or LIA, but overlooks an additional source of unintended output: constitutive transcriptional leakage from the inducible expression cassette. To address this, we used clones expressing only the inducible expression cassettes and found that Gal4UAS-mCMV and all the 6xNFAT variants exhibit basal transcriptional activity in the absence of ligand. These results demonstrate not only that all inducible expression cassettes are functional, but also that transcriptional leakage is a significant contributor to total receptor output in all systems.

An important theoretical advantage of orthogonal systems tested here is that signaling between the synNotch or SNIPR receptors and their inducible promoter, Gal4UAS-mCMV, forms a closed circuit that does not recruit endogenous signaling (Morsut *et al*., 2016; Roybal *et al*., 2016; Manhas *et al*., 2022; Zhu *et al*., 2022). In contrast, TRUCK receptors require that CAR activation is connected to the 6xNFAT signaling by endogenous NFAT signaling (Chmielewski and Abken, 2020), theoretically making the TRUCK systems more prone to leaky transcriptional activity. Surprisingly, the Gal4UAS-mCMV cassette used in synNotch and SNIPR showed higher transcriptional leakage than the 6xNFAT cassettes, calling into question its theoretical advantage.

While the focus on reducing off-target activation is generally receptor ligand-independent activation, our results indicate that refining the transcription regulatory region may significantly reduce off-target activation. For example, given that the 6xNFAT systems were less leaky than Gal4UAS-mCMV, one *a priori* unexpected improvement would be to use a hybrid system. Namely, one in which SNIPR or synNotch are fused to an NFAT-based ATF to activate a 6xNFAT inducible expression cassette.

### TRUCK systems

Preclinical cancer research has established that CAR and TCR-based receptor systems targeting membrane-bound ligands can drive payload production through synthetic NFAT regulatory regions across a range of TRUCK configurations (Chmielewski *et al*., 2011; Chmielewski and Abken, 2017, 2020; Kunert *et al*., 2018; Liu *et al*., 2019; Zimmermann *et al*., 2020; Glienke *et al*., 2022; Harrer *et al*., 2022; Fischer-Riepe *et al*., 2024; Umland *et al*., 2025). However, our findings indicate that receptor activation did not translate efficiently into payload production under the conditions tested. This failure cannot be attributed to a non-functional CAR, as Adu-CAR functionality was confirmed by AβO-induced upregulation of CD69. Further, all 6xNFAT variants exhibited significant transcriptional leakage, indicating active regulatory regions, and all TRUCK variants showed MetLuc induction following PMA/ionomycin stimulation, demonstrating NFAT responsiveness. Nevertheless, no TRUCK variant produced MetLuc significantly above its treatment-matched cassette-only control.

The discrepancy with previous work may be partly explained by the nature of the target antigen and identity of the inducible payload. Unlike the membrane-bound, cancer-associated ligands used in prior TRUCK studies, Adu-CAR in our system targets soluble extracellular aggregates of Aβ, which may engage the CAR differently. Payload identity also appears to be an important determinant of production efficiency in TRUCK systems, independent of receptor activation. Across the literature, single-chain IL-12 variants and IFN are reportedly produced at levels an order of magnitude higher than bioactive IL-18, with the latter falling within the subnanomolar per milliliter range (Chmielewski *et al*., 2011; Chmielewski and Abken, 2017, 2020; Kunert *et al*., 2018; Liu *et al*., 2019; Zimmermann *et al*., 2020; Glienke *et al*., 2022; Harrer *et al*., 2022; Fischer-Riepe *et al*., 2024; Umland *et al*., 2025), suggesting that payload-intrinsic factors are affecting output. In this context, MetLuc production may be constrained by the concurrent expression of chLecanemab from the same inducible cassette, yielding weaker output. Indeed, our results with mIL2 and syn-TRUCK systems do describe a pattern consistent with weak AβO-triggered MetLuc production rather than an outright dysfunctional response.

An anergic-like hyporesponsive state of TRUCK clones might reconcile the presence of a functional Adu-CAR as assessed by the activation marker CD69, and functional expression cassettes, as shown by PMA/Iono treatments, with the lack of a strong MetLuc production in response to AβO treatment. Anergic T cells retain many TCR responses, including upregulation of select activation markers and some effector functions, despite impaired NFAT-driven production of IL-2 (Fathman and Lineberry, 2007). Further, anergic Jurkat and primary T cells retain the capacity to activate NFAT-driven transcription when stimulated with both PMA and ionomycin (Howe *et al*., 2003; Fathman and Lineberry, 2007), mirroring our results showing that PMA/ionomycin rescued MetLuc output across all three TRUCK systems despite the absence of AβO-driven induction.

Anergy can arise from TCR engagement in the absence of sufficient CD28 costimulation (Fathman and Lineberry, 2007), and is one of the reasons second-generation CARs outperform first-generation CD3ζ-only CARs (van der Stegen, Hamieh and Sadelain, 2015). While our Adu-CAR incorporates a CD28 costimulatory domain, we deliberately used CD8α transmembrane regions to minimize the tonic signaling associated with homologous CD28-based constructs (Muller *et al*., 2021). This may have impacted constitutive activation, creating an imbalance between tonic CD28 costimulatory and tonic CD3ζ signaling that predisposes cells to an anergy-like state. If so, substituting the CD8α transmembrane domain with that of CD28 in Adu-CAR may prevent this, albeit at the cost of increased tonic activation. Future research will need to address this possibility.

For now, we show synNotch and SNIPR were clearly superior to all TRUCK variants despite employing the identical Aducanumab scFv as Adu-CAR and the same chLecanemab and MetLuc transgenes used across all 6xNFAT variants.

### SNIPR versus synNotch

We previously demonstrated, for the first time, that a synNotch receptor bearing the same amino acid sequence as the one tested here could drive synthesis and secretion of chLecanemab and chAducanumab in standard IgG and scFv-Fc formats by murine NIH/3T3 cells in response to extracellular Aβ aggregates (Bergo *et al*., 2025). The present results extend this validation to human Jurkat cells, a CD4+ T cell line widely used for *in vitro* CAR characterization (Alonso-Camino *et al*., 2013; Lipowska-Bhalla *et al*., 2013; Bloemberg *et al*., 2020), and further demonstrate that synNotch is clearly superior to all tested TRUCK variants under the conditions examined.

To the best of our knowledge, this is among the first demonstrations of SNIPR responsiveness to extracellular Aβ and the first direct comparison with synNotch in this context, with SNIPR showing superior performance under the tested conditions. However, the distinct contributors to receptor output warrant consideration before interpreting this difference. Both receptors share the same Gal4VP64 ATF, the same downstream Gal4UAS-mCMV regulatory architecture driving chLecanemab and MetLuc transgenes, and an identical Aducanumab scFv. The remaining determinants of receptor output are therefore constitutive off-target activation or LIA, on-target receptor LDA, and antigen availability. Unlike studies in which receptors target an invariant membrane-bound ligand, the AβO preparations used here contain heterogeneous populations of aggregates of varying sizes, which very likely results in greater effective antigen availability for SNIPR than for synNotch. SNIPR has been shown to respond to soluble monomers, whereas synNotch has not (Piraner *et al*., 2025). SynNotch activation requires a minimum mechanical force to elicit activation, a threshold known to require that monomers are immobilized rather than soluble to trigger the receptor (Morsut *et al*., 2016; Lee *et al*., 2023; Brien *et al*., 2024). However, given that both we and others have demonstrated synNotch activation by extracellular proteinaceous aggregates and matrices (Simic *et al*., 2024; Bergo *et al*., 2025; Spetz *et al*., 2025), it is reasonable to infer that aggregates above a certain size also generate the forces required for activation, while smaller assemblies do not. It is therefore likely that lower-order aggregates present in our preparations selectively engage SNIPR but not synNotch, which may alone account for the observed performance difference.

The above notwithstanding, the objective of this comparison is not to evaluate the two receptors under equivalent conditions, but to identify the most suitable platform for AD. From this perspective, the broad size distribution of fibrillar and oligomeric Aβ species that characterizes the AD brain favors a receptor capable of responding across this range, and our results indicate that SNIPR is functionally better suited to this purpose than synNotch.

An unexpected finding was that SNIPR clones did not show the AβO-dependent increase in TMRM signal observed in synNotch and TRUCK clones. Considering TMRM is a mitochondrial dye reflecting both cell viability and activation state in T cells (Ehrenberg *et al*., 1988; Farkas *et al*., 1989; Sukumar *et al*., 2016; Amitrano *et al*., 2021), an activation-dependent increase would be expected in TRUCK cells via CAR-mediated TCR signaling. However, the observation that synNotch clones showed a similar increase suggests that the effect is not strictly dependent on TCR signaling pathways. Moreover, the absence of a comparable increase in expression cassette-only clones suggests that the response requires the presence of an Aβ-binding receptor rather than reflecting a non-specific effect of AβO exposure. One possible explanation is that receptor expression alters the effective presentation of AβO. Because Jurkat cells are a suspension CD4+ T cell line, we speculate that surface-expressed receptors may capture and locally concentrate AβO at the cell surface, whereas in untransduced or cassette-only cells, AβO would be more likely to sediment on the bottom of the well without sustained cell-associated retention. This local enrichment could increase TMRM signal through mechanisms not strictly dependent on the canonical signaling pathway of the Aβ-binding receptor.

As noted, the AβO dependent increase in TMRM for synNotch and TRUCKs appears to be absent in SNIPR-expressing cells. A tentative mechanistic explanation for the divergence of SNIPR relates to its unique mode of activation. Unlike TRUCK and synNotch systems, SNIPR activation by soluble ligands requires endocytic internalization of the receptor-ligand complex (Piraner *et al*., 2025). As a result, AβO may be internalized in SNIPR-expressing cells. Given that Aβ-species have been reported to exert mitochondrial toxicity in other cell types (Walton *et al*., 2020), intracellular accumulation could impair mitochondrial function and offset any activation-dependent increase in TMRM, resulting in a net null effect. However, this remains speculative and warrants investigation in future studies. Regardless, despite this attenuated TMRM response, SNIPR produced the highest payload levels among all systems tested, indicating that its performance is not diminished under these conditions.

Despite all clones receiving identical AβO-FITC concentrations at the onset of the experiment, we found a reduction in Aβ-FITC signal in SNIPR and synNotch groups relative to untransduced cells. Several mechanisms could account for this. Secreted chLecanemab may partially inhibit Aβ aggregate formation, shifting a proportion of Aβ toward more soluble species that are less likely to be retained on the well surfaces through IF washes. Alternatively, surface expression of Aβ-binding SNIPR and synNotch receptors may capture Aβ-FITC on the cell membrane. In that case, unlike untransduced cells, Aβ-FITC-bound SNIPR and synNotch cell clones lost during IF washes may be expected to also result in the loss Aβ-FITC signal in confocal images. Other explanations are possible, and the precise mechanism underlying this apparent clearance will require dedicated future investigation. Nevertheless, despite reduced AβO-FITC retention, both SNIPR and synNotch clones produced chLecanemab in response to AβO-FITC treatment, and the IF results were consistent with those obtained by MetLuc in independent experiments.

Overall, SNIPR outperformed synNotch in our *in vitro* assays and, given its more compact architecture and amenability to humanization, represents a promising receptor platform for targeting Aβ and delivering therapeutic payloads in AD.

### Limitations and Future Directions

Beyond the limitations outlined above, there are additional limitations worth mentioning. All experiments in this study were performed in Jurkat cells, which are a CAR-competent cell line to compare TRUCKs, synNotch and SNIPR. TRUCKs may behave differently in primary T cells and other CD3 signaling cells. However, synNotch and SNIPR are orthogonal and should work in any cell type, they may be variations in their behavior in different cell types. Thus, future studies should evaluate the system directly in the intended therapeutic cell types.

This study indicates that off-target signaling arises from both receptor LIA and the TL from the inducible expression cassette. Future work should therefore address both the receptor and inducible expression cassette components to minimize off-target signaling. A complementary strategy at the payload level may also be valuable to limit the consequences of off-target secretion. For example, in conventional antibody therapies, neonatal Fc receptor (FcRn) epitopes are incorporated to extend systemic half-life (Pyzik *et al*., 2023), but in cell-based CNS delivery the opposite approach may be advantageous: engineering payloads such as Lecanemab for reduced systemic stability would allow rapid clearance of any off-target secretion in the blood.

A notable advantage of TRUCK systems is that the CAR simultaneously mediates T cell enrichment at the target site and drives payload delivery from a single receptor. In contrast, synNotch and SNIPR systems require the addition of a separate CAR to achieve targeted enrichment, increasing the size and complexity of the constructs required. Furthermore, certain SNIPR and CAR configurations have been shown to interact (Zhu *et al*., 2022), making it important for future studies to assess SNIPR and CAR receptors in tandem to fully characterize their combined behavior.

In cell-based delivery, where cells can self-renew and potentially produce payload indefinitely, effective dose can be modulated indirectly through the number of adoptively transferred cells and by limiting persistence via suicide genes (Sahillioglu and Schumacher, 2022). The systems described here offer an additional, self-limiting mode of regulation: receptor engagement with Aβ aggregates drives secretion of an Aβ-clearing antibody that eliminates its own activating stimulus, thereby shutting off further production. This closed-loop architecture operates under a fundamentally different framework from conventional drug administration, in which a finite dose is delivered at a defined concentration. Establishing the conceptual and practical standards for dosimetry in cell-based therapeutic delivery will be an important challenge for the field going forward.

## Conclusions

This study provides the first direct comparison of three TRUCK systems with synNotch and SNIPR for Aβ-targeted inducible payload delivery under matched conditions. Both orthogonal systems substantially outperformed TRUCKs, not because they exhibited tighter basal control, but because the TRUCK systems failed to translate Aβ-induced receptor activation into robust inducible payload production. Among orthogonal platforms, SNIPR consistently exceeded synNotch, an advantage likely rooted in its documented capacity to respond to soluble proteins (Piraner *et al*., 2025), making it well suited to the heterogeneous Aβ landscape of the AD brain.

## Acknowledgements

We thank Akos Gerencser for conceptual guidance on the Revvity G3 Explorer platform and experimental design. The figures were prepared using BioRender.

## Declarations

### Author Contributions

CJS and CCW conceptualized the study and designed the constructs. Subcloning was performed by CJS, CCW, and ZM. Lentivirus production and generation of cell clones were carried out by CJS. CJS and CCW designed the experiments. Automated workflow with high-content imaging experiments were conducted by CJS and IB, and all other experiments were performed by CJS. Data processing and statistical analyses were carried out by CJS and CCW. JKA and CCW supervised the project and provided overall guidance. CJS and CCW wrote the manuscript with input from all authors.

### Data Availability

Datasets for the current study are available from the corresponding author on reasonable request

### Ethics approval

This article does not contain any studies with human or animal subjects performed by any of the authors.

### Consent to participate

The authors declare their agreement to participate.

### Code availability

Not applicable.

### Artificial intelligence (AI)

Not applicable.

