## Supplementary Materials for "*In vitro* comparison of Aβ-targeting SNIPR, synNotch, and TRUCK for cell-based drug delivery in Alzheimer’s disease"

**Methods**

**Constructs**

All constructs were incorporated into self-inactivating lentiviral transfer vectors to prevent transcriptional readthrough of the inducible gene from the long terminal repeats following integration. The generation of the TRUCK, synNotch, and SNIPR receptor systems, as well as the expression cassette-only constructs, are described in the Results and Supplementary Materials. Relevant amino acid and nucleotide sequences are listed in Table S1. All constructs were codon-optimized for expression in human cells, synthesized, subcloned, and sequence-verified by VectorBuilder.

**Lentiviral production**

The human embryonic kidney (HEK) 293T/17 adherent cell line (ATCC®, CRL-11268™) was used for lentiviral production. The cells were maintained in Dulbecco's Modified Eagle's Medium (DMEM; VWR, 45000-304) supplemented with 10% Fetal Bovine Serum (FBS; VWR, 45000-734), 100 U/mL penicillin and 100 µg/mL streptomycin (1X Pen/Strep; Gibco, CAT# 15140122) at 37°C with 5% CO₂, and switched to antibiotic-free medium prior to transfection. All constructs were delivered using third-generation lentiviral vectors (Dull *et al.*, 1998). Lentiviral particles were produced by co-transfecting HEK293T/17 cells with lentiviral transfer plasmids and the third-generation packaging plasmids pMDLg/pRRE (Addgene #12251; RRID:Addgene_12251), pRSV-Rev (Addgene #12253; RRID:Addgene_12253), and pMD2.G (Addgene #12259; RRID:Addgene_12259). pMDLg/pRRE, pRSV-Rev, and pMD2.G were gifts from Didier Trono (École Polytechnique Fédérale de Lausanne, EPFL). Following the manufacturer's instructions, lentiviral plasmids were transfected into HEK293T/17 cells using Lipofectamine™ 3000 Transfection Reagent (Invitrogen, CAT# L3000001). Viral media was replaced with antibiotic-free complete media 6 h post-transfection. Lentiviral supernatants were harvested 24 h and 48-52 h post-transfection, centrifuged at 800 × g for 10 min at 4°C, filtered through a 0.2 µm PES filter (Fisher Scientific, CAT# 50-104-9880), and subsequently frozen as single-use aliquots at −80°C for long-term storage.

**Lentiviral transduction**

**Single-vector TRUCK systems:** Cells were seeded in antibiotic-free complete medium in 12-well plates at 7.5 x 10⁵ cells per well. Polybrene was added to a final concentration of 8 µg/mL (VectorBuilder, CAT# PL0001) with the lentiviral particles in a final volume of 1 mL per well. The plates were wrapped in plastic and centrifuged at 1,000 x g for 60 min at 32 °C, followed by incubation at 37 °C with 5% CO₂ for 16-18 h. The viral particle-containing media were replaced with fresh complete culture media, and the transduced cells were purified by FACS 2 days after media replacement. **Two-vector systems:** Cells were first transduced with the inducible expression cassettes using the polybrene spinoculation protocol described above. Transduced cells were sorted by FACS one day after replacing the viral particle-containing media and transduced a second time with receptor vectors, either using the polybrene spinoculation protocol described above or the Retronectin-virus-bound method (RBV) with spinoculation. For RBV, untreated 12-well plates were coated with 1 mL of 24.5 µg/mL Retronectin (Takara, CAT# T100B) in Dulbecco’s Phosphate Buffered Saline 1X (DPBS; VWR, CAT# 02–01119-0500) overnight at 4 °C, blocked with 2% BSA in DPBS for 30 min, and washed twice with DPBS. Only the 8 central wells of the 12-well plates were used, as the 4 wells in the corners crack during centrifugation. Receptor-vector lentiviral particles were diluted in antibiotic-free complete medium to reach a final volume of 1 mL per well. The plates were wrapped in plastic for safety and centrifuged at 1,500 x g for 2 h at 32°C to allow virus binding. Wells were washed once with 2% BSA in DPBS, and cells were seeded at 1 x 10⁵ cells/mL in 1 mL per well. Plates were wrapped again, centrifuged at 800 × g for 30 min at 32°C, and incubated at 37°C with 5% CO₂ for 1-2 days before sorting.

**Cell culture**

**Expansion:** The Jurkat Clone E6-1 cell line (ATCC®, TIB-152™) was cultured in suspension using untreated plates or flasks at 37°C with 5% CO₂. Cells were cultured in complete culture media comprising Roswell Park Memorial Institute (RPMI) 1640 Medium (Corning, CAT# 10-040-CV), 10% heat-inactivated Fetal Bovine Serum (hiFBS; Corning, CAT# 35-011-CV), and 1X Pen/Strep. Cells were cryopreserved in freezing medium (RPMI, 10% hiFBS, 5% Dimethyl Sulfoxide (DMSO; Sigma Aldrich, CAT# D2650). **Single-vector TRUCK systems CD69 and Metluc experiments:** Cells were seeded in uncoated 96-well plates to obtain a final assay density of 2 x 10^5^ cells/mL. Cells were then treated with AβO or vehicle for the total 46 h assay duration or with 25 ng/mL phorbol 12-myristate 13-acetate (PMA; Sigma-Aldrich, CAT# P1585) and 1000 ng/mL ionomycin (Iono; Sigma-Aldrich, CAT# I3909) for the last 6 hours. At the end of the assay, cells and media were collected for analysis by flow cytometry and Metluc, respectively. **TRUCKs, synNotch, and SNIPR experiments:** For the MetLuc assay and high-content TMRM and EGFP imaging, a glass-bottom 384-well plate (Cellvis, CAT# P384-1.5H-N) was coated with 50 µg/mL Poly-D-Lysine (PDL; Gibco, CAT# A3890401), sealed, and incubated at 4°C overnight. The wells were washed 3 times for 1 h with sterile Milli-Q water, then once with complete culture media. Cells were then plated at 6,000 cells per well and subsequently treated with AβO or vehicle for 48 h in a final volume of 60 µL per well. At the end of the assay, live cells were imaged for TMRM and EGFP, while the media was collected for MetLuc analysis. **SynNotch and SNIPR ICC experiments:** µ-Slide 8 Well^high^ ibiTreat chambers (Ibidi, CAT# 80806) were coated with 50 µg/mL PDL as described above. Cells were subsequently plated at 80,000 cells per well and treated with AβO-FITC for 48 h in a final volume of 200 µL per well, after which the cell cultures were fixed for ICC.

**Amyloid-β oligomer preparations**

Beta amyloid (Aβ; 1-42) aggregation kit (rPeptide, A-1170-2) and lyophilized FITC-labeled Aβ (1-42) peptides (rPeptide, A-1119r-2) were reconstituted in 5 mM Tris at a concentration of 1 mg/mL and diluted with HPLC water and Tris-buffered saline (TBS) to obtain a 100 µM stock solution. Aβ oligomer-enriched preparations were generated as previously described (Bergo *et al.*, 2025; Siebrand *et al.*, 2025). Stock solutions were incubated at 37°C for 3 h. Vehicle solutions consisted of equivalent volumes of Tris, HPLC water, and TBS, incubated under the same conditions.

**Flow Cytometry**

Cells were collected from cultures, washed with DPBS, and stained with the LIVE/DEAD^™^ Fixable Violet Dead Cell Stain Kit (1:1000, Invitrogen, CAT# L34964) for 20-30 min on ice. Cells were washed again with DPBS and stained for 20-30 min on ice with HA Epitope Tag Polyclonal Antibody, DyLight™ 549 (1:1000, ThermoFisher, CAT# 600-442-384). For flow cytometric analysis of activation, anti-human CD69-FITC (1:400, FN50, BioLegend, CAT# 310904) was added to the staining panel. Cells were then washed in DPBS and resuspended in FACS buffer (DPBS with 1% FBS and 10 mM HEPES; Gibco, CAT# 15630080). Transduced cells were sorted based on expression of HA-tag, EGFP, or both, using a BD FACSAria II cell sorter (BD Biosciences). Viability, HA-tag, and CD69 expression were measured using a Cytek Aurora (Cytek). Gates were created with FlowJo™ Software (RRID:SCR_008520) and exported for statistical analyses.

**MetLuc assay**

MetLuc reporter activity was measured using the Ready-To-Glow™ Secreted Luciferase Reporter Assay (Clontech, CAT# 631728) according to the manufacturer’s instructions in a white, flat-bottom LUMITRAC™ 96-well plate (Greiner Bio-One, CAT# 655074). Luminescence was measured using a BioTek Cytation3 multimode plate reader and reported as relative light units (RLU).

**Automation, liquid handling, and high-content imaging**

Automated workflow execution, including plate handling and instrument operations, was implemented using plate::works™ (Revvity) software. At the end of the treatments, a plate::handler™ FLEX (Revvity) transferred the 384-well plate with live cells to a Microplate Centrifuge with Automated Centrifuge Loader (Agilent) for centrifugation at 100 x g for 1 min. Supernatant was separated from cells using the JANUS™ G3 Automated Liquid Handling Workstation (Revvity) equipped with a 96-channel Modular Dispense Technology™ (MDT) dispense head (Revvity), programmed to uniformly transfer 50 µL of supernatant from each well of the 384-well plate for the MetLuc assay, while leaving 10 µL per well containing the centrifuged cells. An Echo® 650 Series Liquid Handler (Beckman Coulter) then dispensed TMRM at a final concentration of 7.5 nM TMRM into the remaining well volume. This plate with cells was then incubated for 30 min at 37°C with 5% CO₂ in a LiCONiC STX44 Automated Incubator, after which it was shuttled by the plate::handler™ FLEX to the Operetta CLS High-Content Analysis System (Revvity) to image the live cells. Four images per well were acquired by the Operetta CLS using a 20x objective (NA 0.4) to obtain TMRM and EGFP fluorescence signals, which were used for subsequent image analysis.

**Immunofluorescence and microscopy**

Immunostaining and confocal imaging were performed without removing the cells from the µ-Slide 8-well chambers. Wells were fixed for 6 min at room temperature (RT) with 2% paraformaldehyde (PFA; Santa Cruz, 3052589-4), followed by two 6-min washes with ice-cold PBS. Residual PFA was quenched by a 15 min incubation with 50 mM NH₄Cl in PBS, followed by a 6 min wash in PBS, and a 30 min RT incubation with 10% normal goat serum (NGS; Jackson ImmunoResearch, 005-000-121) in PBS to block non-specific antibody binding. Because immunocytochemistry was performed to detect extracellular proteins, no permeabilization step was included. After an additional wash in PBS, the IgG2a Fc domain of chLecanemab was detected by incubating the wells for 1 h at RT with anti-mouse IgG2a–Alexa Fluor 546 (1:500; Invitrogen, A-21133) diluted in PBS containing 1% NGS. Wells were then washed two times for 10 min with PBS containing 0.5% Tween-20 (PBST). Nuclear staining was performed with PBS containing 1 µg/mL 4′,6-diamidino-2-phenylindole (DAPI; Invitrogen, D1306; 1:1,000 from a 1 mg/mL stock). After two additional PBS washes, PBS was fully replaced with glycerol (Invitrogen, 15514-011). Confocal tile-scan images were acquired using a Zeiss LSM 980 confocal microscope with a 20x objective.

**Image Processing**

**TMRM and EGFP assays**: The 2D images obtained from the Operetta CLS were processed using the software Harmony 5.2.3 with PhenoLOGIC™. ROI were generated using the Find Cells function in the Digital Phase Contrast channel. Morphology and Intensity properties were calculated, and supervised classification was applied using the Linear Classifier function to select ROIs representing valid cells while minimizing artifact selection. Pixel intensities of TMRM and EGFP within the ROIs were then exported for statistical analyses. The image processing workflow is also described in Sup. Fig. 2f. **Aβ-FITC and chLecanemab analysis:** Images were generated using ZEISS ZEN Microscopy Software (RRID:SCR_013672) and IMARIS software (RRID:SCR_007370). Image processing was performed on 3D images using IMARIS software as described in Sup. Fig. 4d, e. Voxel (volumetric pixel) signal intensities captured by the 3D ROIs were exported in CSV files for statistical analysis.

**Statistical analysis**

Statistical analyses were performed using JASP (RRID:SCR_015823) and the built-in statistical tools in BioRender (BioRender.com), the latter powered by R (RRID:SCR_001905). Outliers were identified using box plots, normality was assessed with the Shapiro-Wilk test (p > .05) and Q-Q plots, and homogeneity of variances was evaluated using Levene's test on the mean (p > .05) and predicted vs. residual scatter plots. One-way or two-way ANOVAs were applied when parametric assumptions were met. When assumptions were violated, appropriate data transformations were applied based on skewness and residual plots, and assumptions were retested. For persistent violations of homogeneity of variances in two-way ANOVAs, groups were analyzed by a one-way Welch's F-test. There were no violations of normality requiring non-parametric tests. Post-hoc comparisons used Tukey all-pairwise or Bonferroni-corrected planned comparisons when assumptions were met, and Games-Howell comparisons when homogeneity of variances was violated. In the presence of a non-significant omnibus test, exploratory post-hoc comparisons were performed to identify potentially biologically relevant effects. Exact p-values are reported in Tables S2–5.

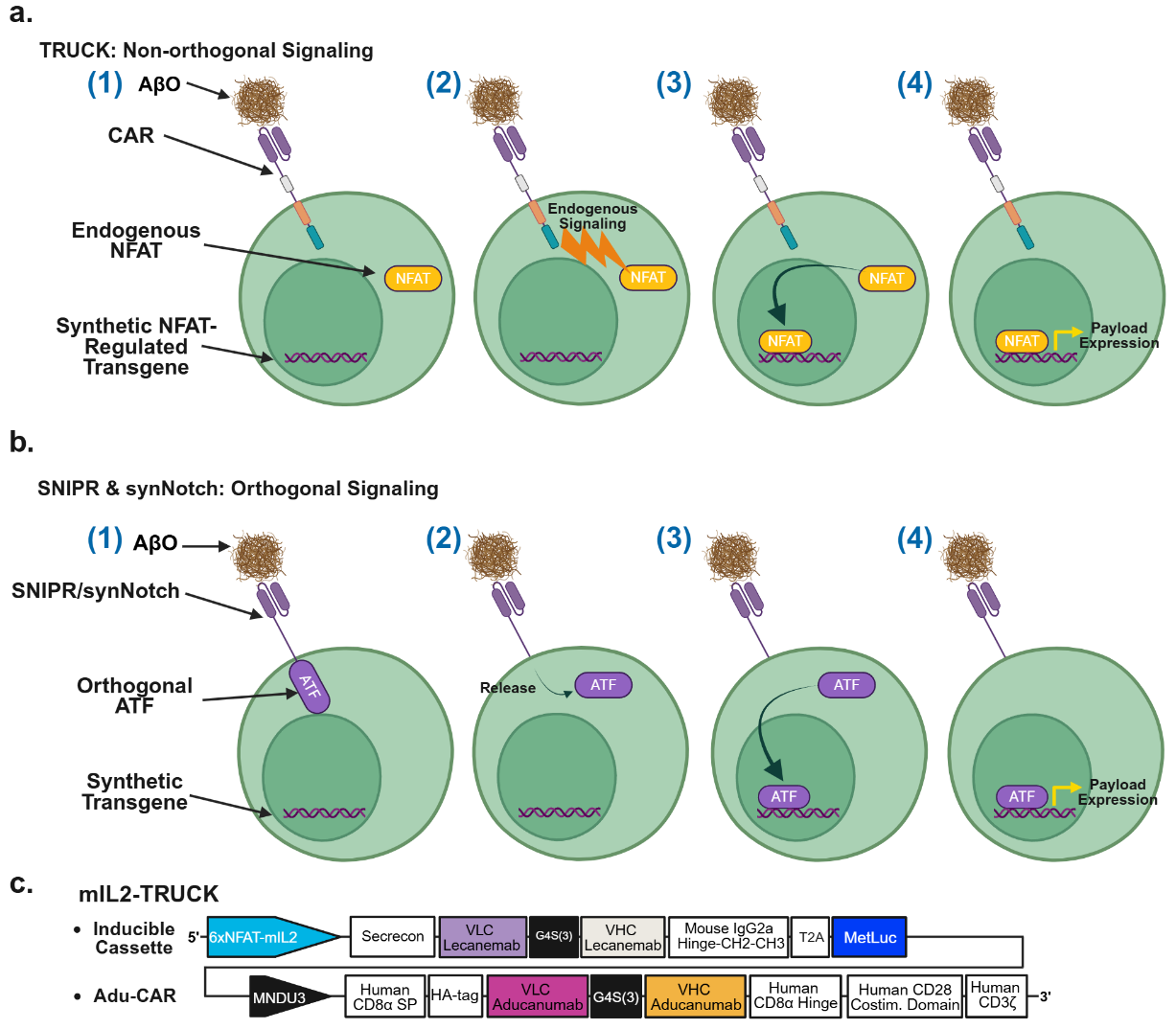

**Supplementary Figures 1a-c.** **a**. **TRUCK: Non-orthogonal signaling**. (**1**) An Aβ-targeting CAR regulates transgene expression through endogenous NFAT signaling. (**2**) Antigen (AβO) engagement activates CAR signaling, leading to downstream TCR-like signaling cascades. (**3**) This results in NFAT dephosphorylation and nuclear translocation. (**4**) Nuclear NFAT binds NFAT-responsive elements upstream of a minimal promoter, inducing transgene expression. **b**. **SNIPR and synNotch: Orthogonal signaling.** (**1**) Aβ-targeting synNotch or SNIPR receptors regulate transgene expression via a receptor-tethered artificial transcription factor (**ATF**). (**2**) Antigen (AβO) binding triggers the proteolytic cleavage within the transmembrane domain of the receptors and the release of the intracellular ATF. (**3**) Nuclear localization signals (NLS) direct the ATF to the nucleus. (**4**) The ATF binds its cognate response elements to initiate transcription of the transgene independently of endogenous signaling pathways. **c**. Schematic representation of the Aβ-targeting mIL2-TRUCK construct, comprising a 6×NFAT–mIL2 inducible expression cassette followed by a constitutively expressed Adu-CAR. The inducible expression cassette contains a chimeric human-mouse chLecanemab, directed for secretion by a secrecon signal, with the variable light and heavy chains (VLC & VHC) of Lecanemab fused to the hinge of mouse IgG2a as well as an in frame Metluc separated by a T2A sequence. Relevant nucleotide and amino acid sequences are provided in Table S1a–c.

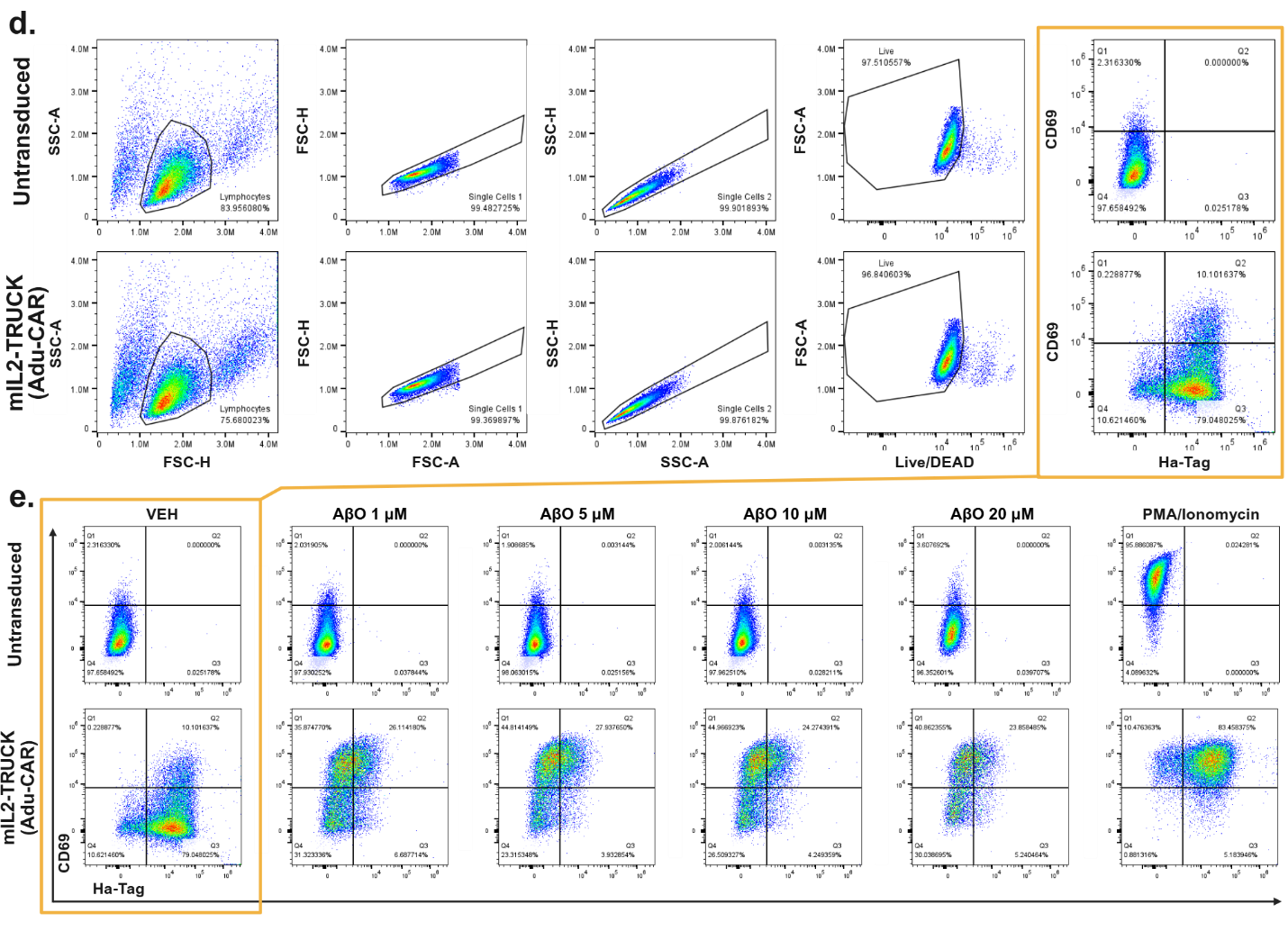

**Supplementary Figures 1d, e**. **d**. Representative density plots showing Live/Dead Fixable Violet staining (viability), CD69 expression (activation), and HA-tag immunostaining (CAR surface abundance) in VEH-treated untransduced (UT) and mIL2-TRUCK–expressing Jurkat cells. **e**. Representative CD69 and HA-tag plots for UT and mIL2-TRUCK cells under the indicated VEH, AβO, and PMA/ionomycin (PMA/Iono) treatments. HA-tag signal is reduced under AβO treatment relative to VEH and PMA/Iono conditions, consistent with receptor internalization and/or steric masking of the HA epitope upon antigen engagement.

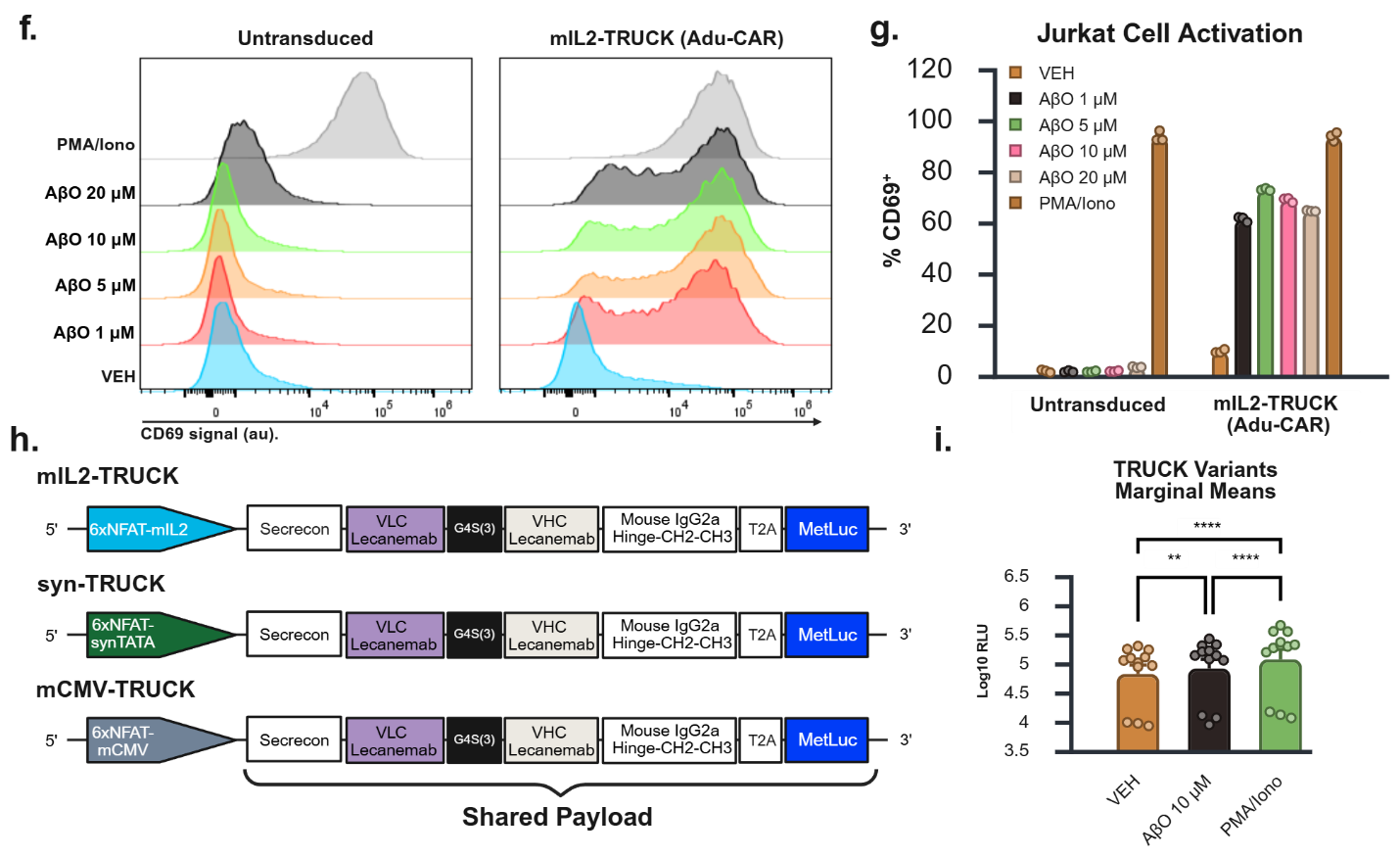

**Supplementary Figures 1f-i**. **f**. Representative flow cytometry histograms showing CD69 fluorescence intensity (arbitrary units, au) in untransduced (UT) and Adu-CAR–expressing mIL2-TRUCK Jurkat clones under the indicated treatment conditions. PMA/ionomycin (PMA/Iono) induced CD69 expression in both UT and transduced cells. In contrast, increased CD69 signal in response to AβO was observed only in Adu-CAR–expressing clones. **g**. Percentage of CD69-positive (CD69⁺) cells in UT and mIL2-TRUCK (Adu-CAR) clones. **h**. Schematic representation of the 6xNFAT inducible expression cassettes used in the mIL2-, synTATA-, and mCMV-TRUCK systems. The constructs are identical except for the minimal promoter region (mIL2, synTATA, or mCMV). Relevant nucleotide sequences are provided in Table S1a, d, and e. **i**. Treatment marginal means of MetLuc production across VEH, AβO, and PMA/Iono conditions. Treatment marginal means represent the average MetLuc output across all clones within each treatment. The lower datapoints within each treatment correspond to MetLuc production by untransduced (UT) cells. Statistical analyses: (i) Bonferroni corrected all pairwise comparisons. Full statistical analysis available in Table S2d. ** p < 0.01, *** p < 0.001, **** p < 0.0001. Data are presented as mean ± SEM from at least three biological replicates per treatment-by-clone combination.

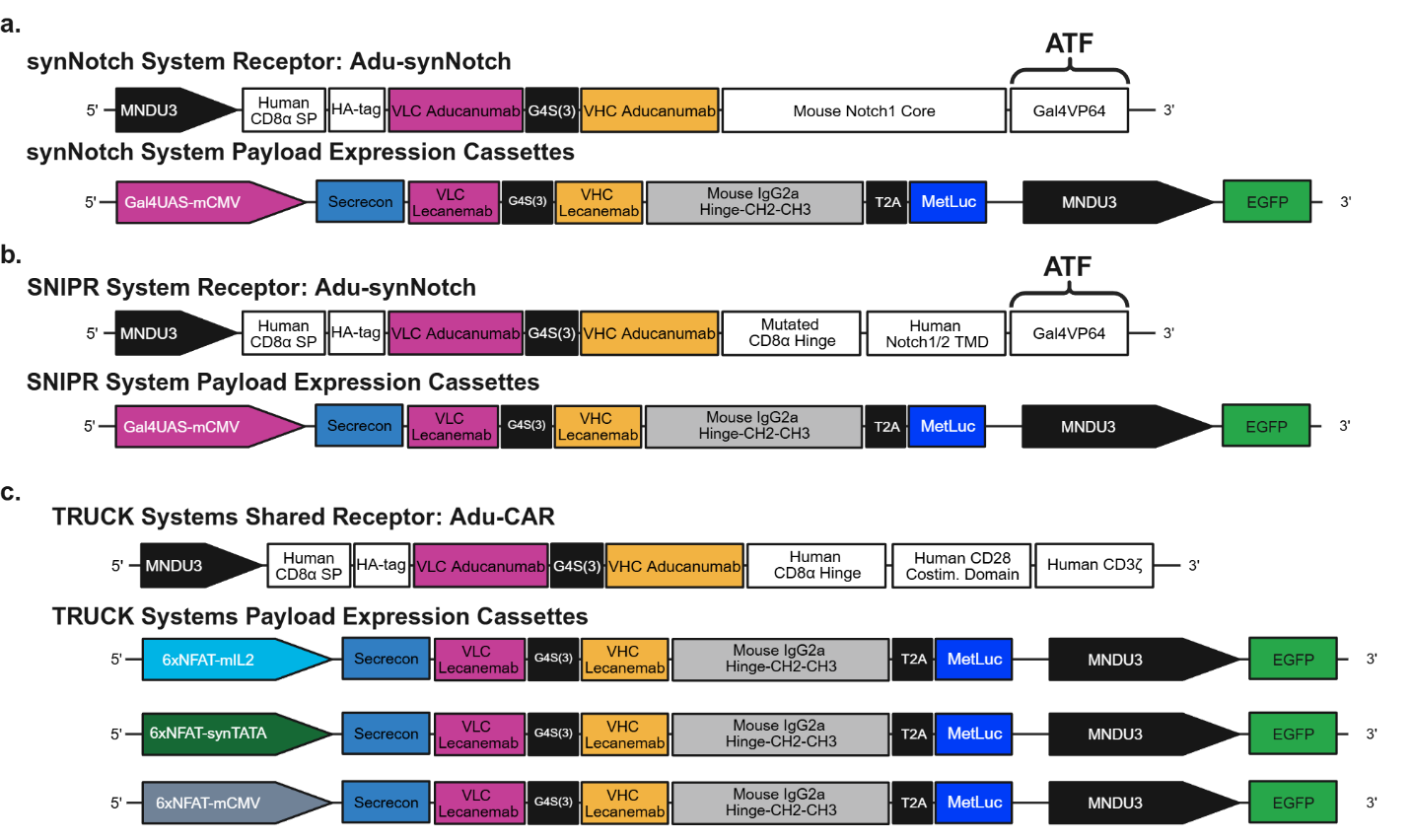

**Supplementary Figures 2a-c.** **a.** Architecture of Aβ-targeting synNotch, SNIPR, and TRUCK systems. **a–c**. Schematic representation of the two-vector design used for (**a**) synNotch, (**b**) SNIPR, and (**c**) TRUCK systems. In all cases, one vector encodes the receptor and the second vector encodes the inducible payload expression cassette. All receptor constructs contain a CD8α signal peptide (SP), an N-terminal HA-tag, and an Aducanumab-derived scFv composed of variable light (VLC) and variable heavy (VHC) domains. **a.** synNotch receptor incorporating the murine Notch1 core regulatory domain fused to a Gal4VP64 artificial transcription factor (**ATF**). **b.** SNIPR receptor containing a modified CD8α hinge region to enhance activation and human Notch1/Notch2 transmembrane domains fused to the same Gal4VP64 ATF used in synNotch. The inducible expression cassette is identical between synNotch and SNIPR systems. **c**. TRUCK variants use the same Adu-CAR receptor vector. The inducible expression cassettes are delivered on a separate vector and differ only in the minimal promoter region (mIL2, synTATA, or mCMV). Across all platforms, the inducible expression cassette encodes a chLecanemab using a secrecon signal sequence for secretion and secreted MetLuc separated by a T2A sequence, followed by a separate MNDU3 promoter constitutively expressing an EGFP reporter. With this two-vector system, receptor-transduced Jurkat cells are detected by the HA-tag, and transduction with the inducible expression cassette vector is detected by the constitutively expressed EGFP. Relevant nucleotide and amino acid sequences are provided in Table S1a–h.

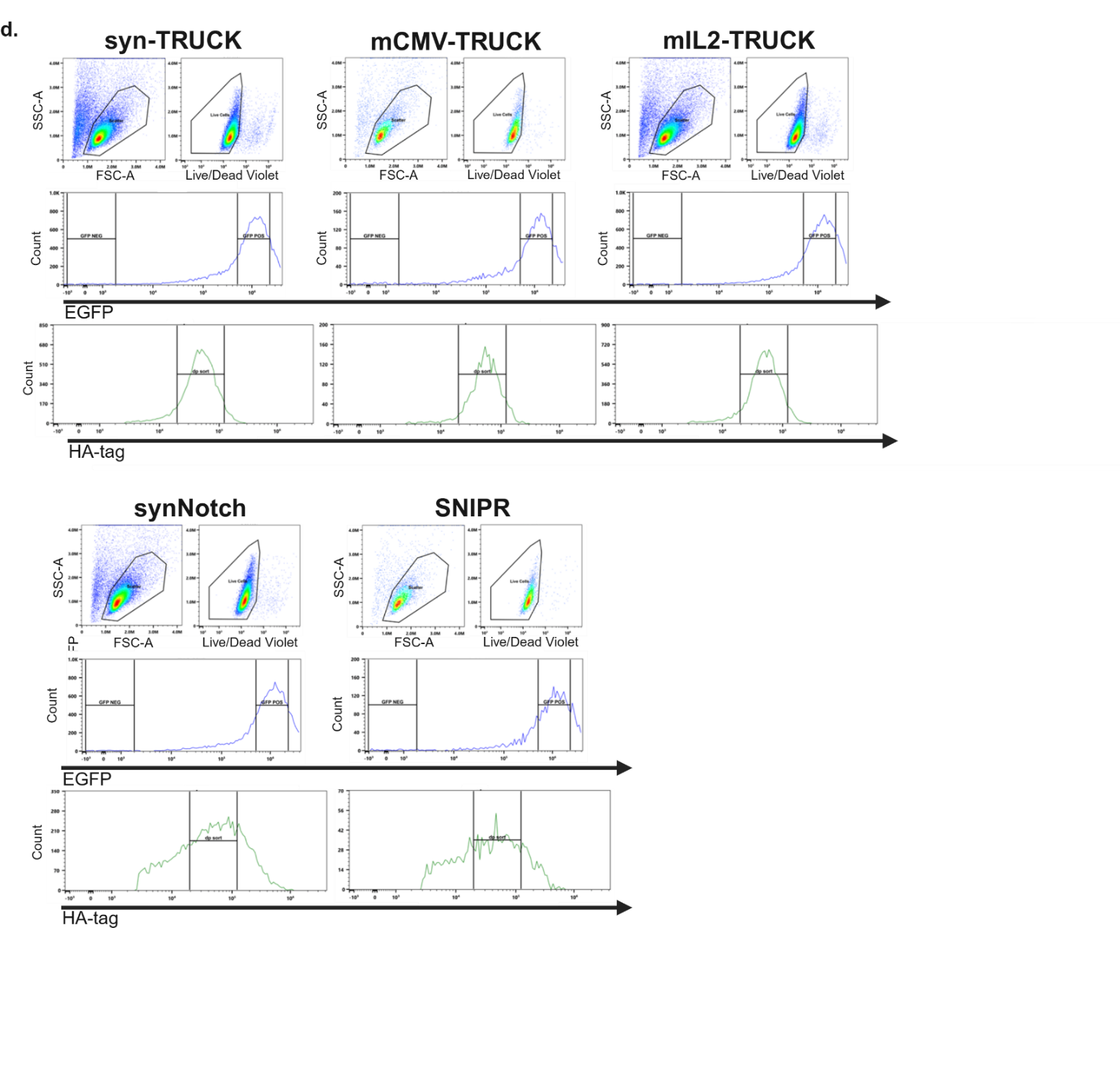

**Supplementary Figure 2d**. Representative flow cytometry plots illustrating the gating strategy used for fluorescence-activated cell sorting (FACS) of cells expressing the complete receptor systems. Live/Dead Fixable Violet staining was used to exclude non-viable cells. Subsequently, a standardized EGFP fluorescence intensity range was first applied to select cells expressing the inducible payload cassette (right gate), ensuring comparable cassette expression across clones. Within this population, a standardized HA-tag intensity range was applied to select receptor-expressing cells, ensuring homogeneous constitutive receptor expression.

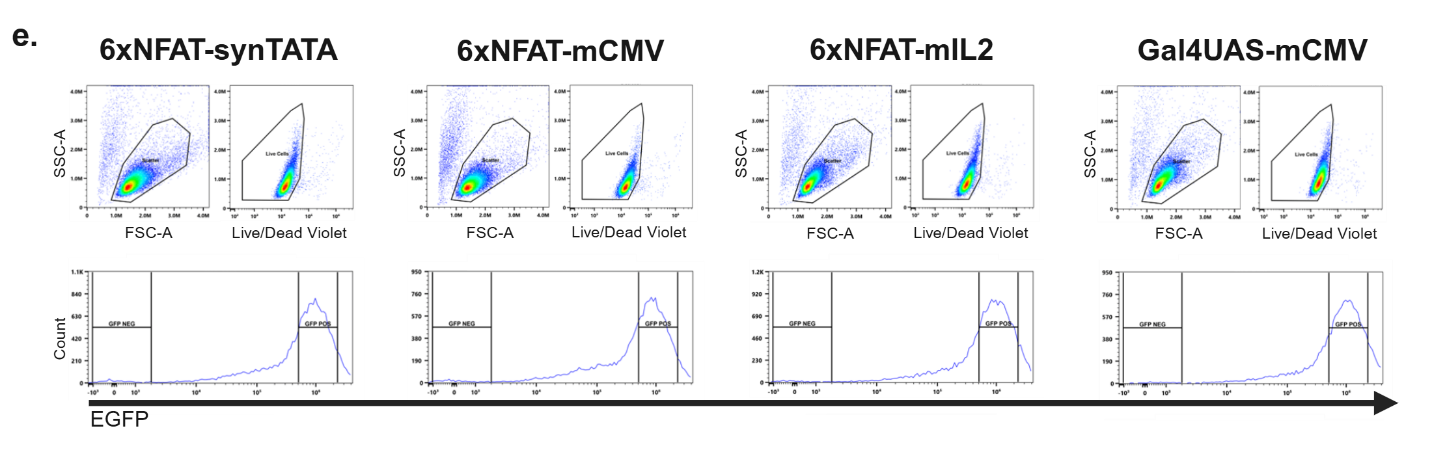

**Supplementary Figure 2e**. Representative flow cytometry plots illustrating the gating strategy used for fluorescence-activated cell sorting (FACS) of cells expressing only the inducible expression cassettes. Live/Dead Fixable Violet staining was used to exclude non-viable cells. Subsequently, the same standardized EGFP fluorescence intensity range as used for the full receptor systems (Sup. Fig. 2d) was applied to select cells expressing the inducible payload cassette (right gate) to ensure comparable expression across clones.

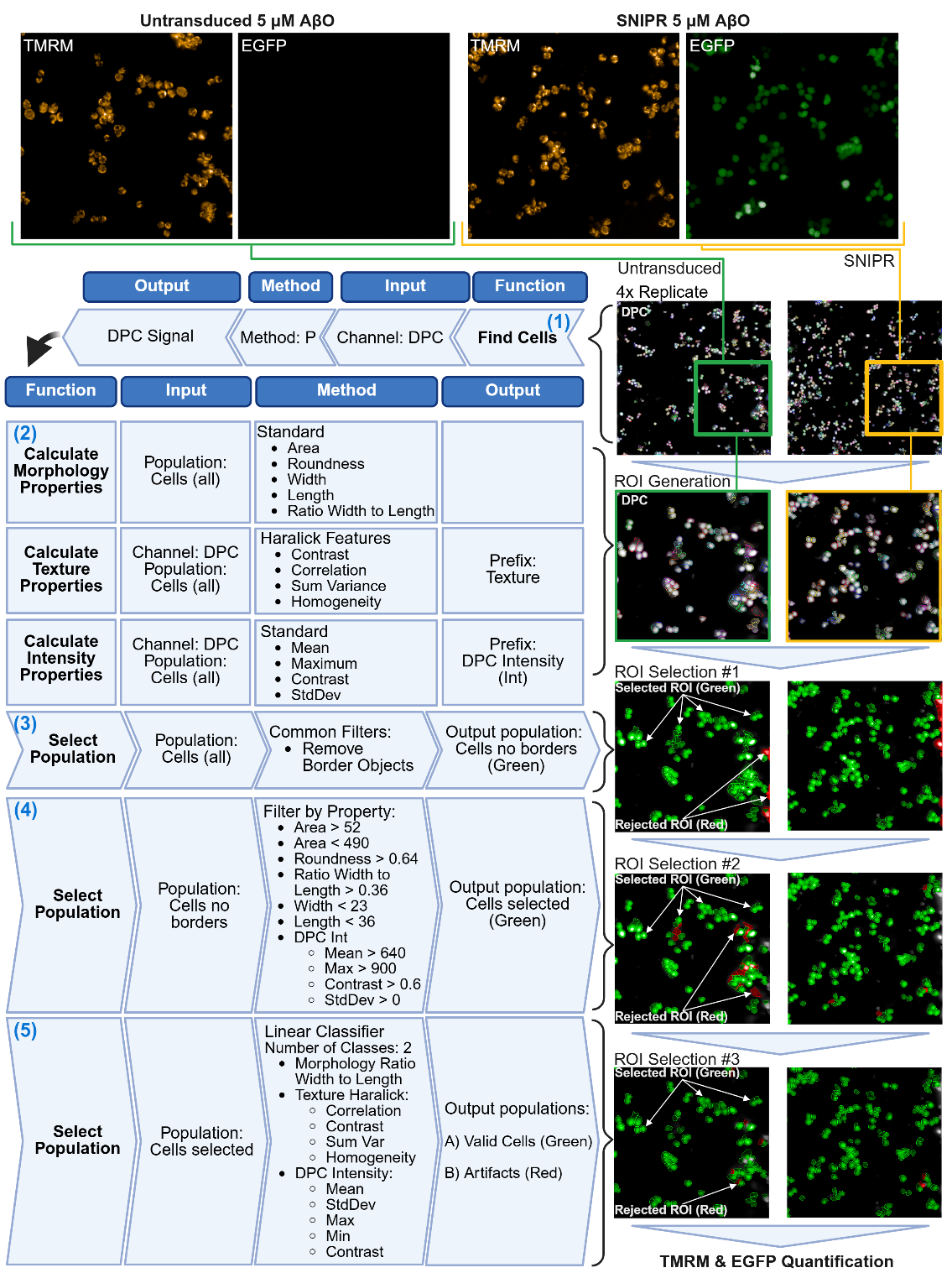

**Supplementary Figure 2f**. Workflow for analysis of Digital Phase Contrast (DPC), TMRM, and EGFP microscopy images using Harmony 5.2 with PhenoLOGIC™. Representative TMRM and EGFP images of untransduced and SNIPR-transduced Jurkat cells treated with 5 μM ABO are shown (top). Insets show expanded views of selected regions (green and orange squares, top right) for increased visual detail. Four images per treatment per replicate were acquired for analysis. **(1)** The Find Cells function segments DPC signal to distinguish cells from background. **(2)** Morphological, Texture, and Intensity Properties are calculated using Standard and Haralick Features to generate ROIs around individual cells. **(3–5)** Sequential filtering steps retain valid ROIs (green) while progressively removing artifactual ROIs (red). Specifically, the Select Population function with user-defined criteria excludes cells that intersect image borders (**3**) and cells that fail morphological or signal-intensity thresholds (**4**). A machine learning Linear Classifier is then trained to identify the final set of valid ROIs (**5**), which are used for quantification of TMRM and EGFP signal intensities. A total of 134-688 objects (ROIs), each representing a single cell, were analyzed per biological replicate. Images were captured using Operetta high-content CLS with a 20x objective.

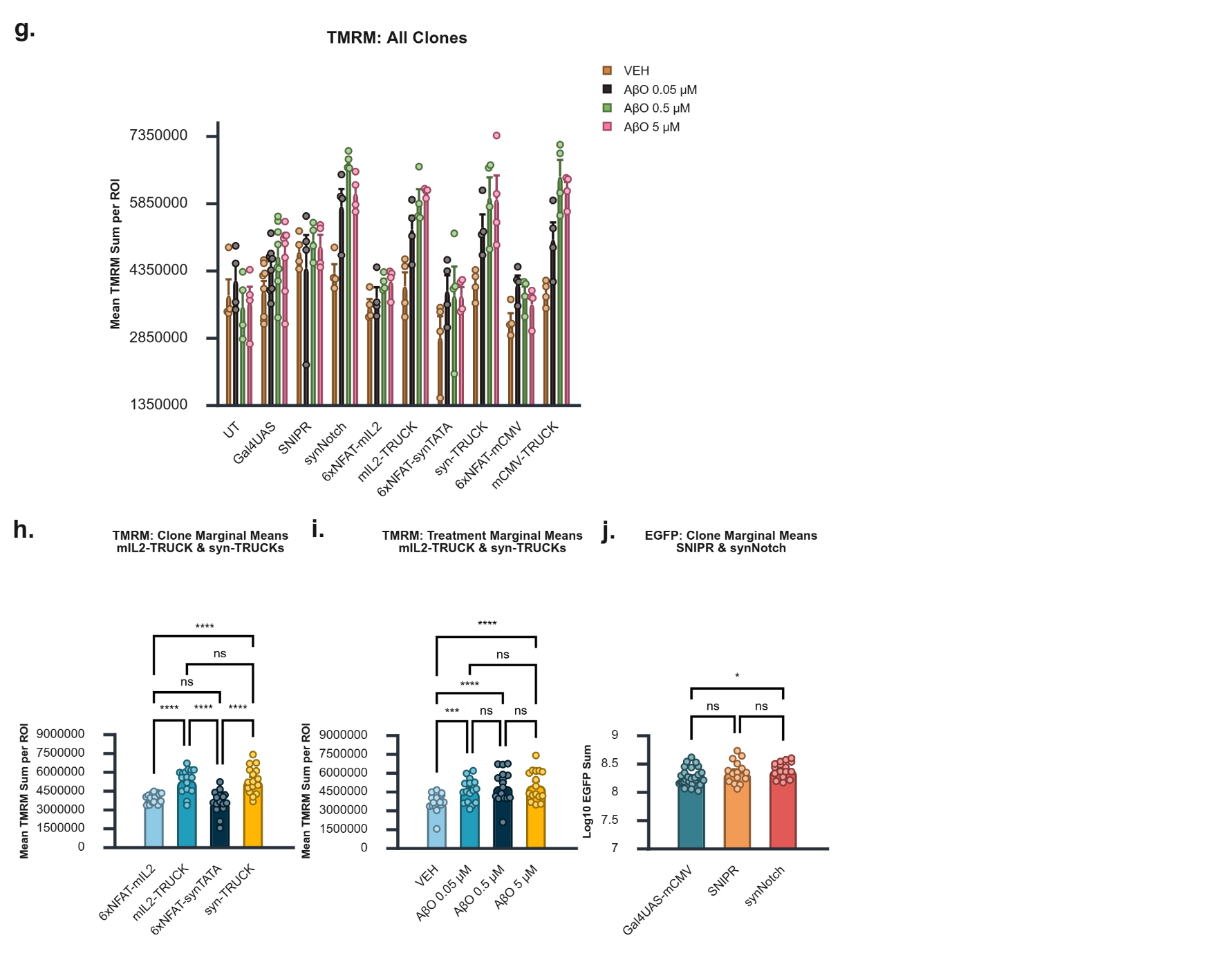

**Supplementary Figure 2g-j. g.** Mean TMRM signal sum per region of interest (ROI) across all experimental groups. This metric serves as a proxy for the average total TMRM fluorescence per cell. Statistical analyses were performed separately for three system categories: synNotch and SNIPR (Fig. 2b), mIL2- and syn-TRUCKs (Fig. 2c), and mCMV-TRUCKs (Fig. 2d). **h, i.** Marginal mean multiple comparisons of mean TMRM signal sum for the mIL2- and syn-TRUCK systems, shown for **(h)** clone and **(i)** treatment main effects. Clone-by-treatment interaction comparisons are presented in Fig. 2c. Full statistical analysis available in Table S3b**. j.** Marginal mean multiple comparisons of Log10-transformed EGFP signal sum for clone main effects within the synNotch and SNIPR systems. The EGFP sum aggregates the total EGFP fluorescence across all ROIs per well and serves as a proxy for cumulative expression of the inducible expression cassette. Clone by treatment multiple comparisons in Sup. Fig. 2k. Full statistical analysis available in Table S3d. Statistical analyses: (**h-j**) Bonferroni corrected all pairwise multiple comparisons. * p < 0.05, ** p < 0.01, *** p < 0.001, **** p < 0.0001. Data are presented as mean ± SEM from at least three biological replicates.

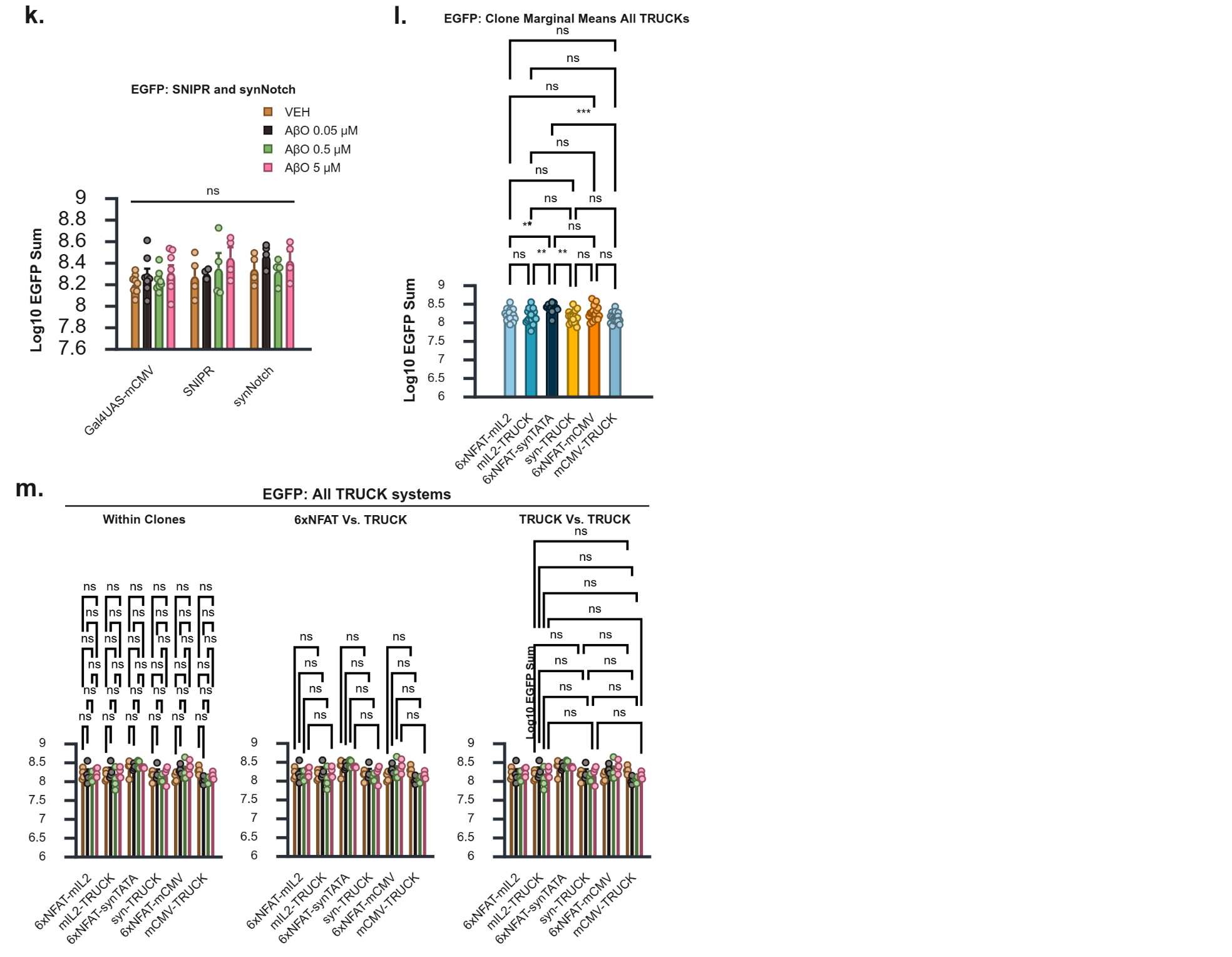

**Supplementary Figure 2k-m. k.** Planned comparisons of Log10-transformed EGFP signal sum by treatment group for the SNIPR and synNotch systems. Full statistical analysis available in Table S3d.  **l.** Marginal mean multiple comparisons of Log10-transformed EGFP signal sum for clone main effects within the mIL2-, syn-, and mCMV-TRUCK systems. Clone by treatment multiple comparisons in Sup. Fig. 2m. Full statistical analysis available in Table S3e. **m.** Planned comparisons of Log10-transformed EGFP signal sum by treatment group for the mIL2-, syn-, and mCMV-TRUCK systems. Full statistical analysis available in Table S3e. Statistical analyses: Bonferroni corrected (**l**) all pairwise and (**k,** **m**) planned multiple comparisons. * p < 0.05, ** p < 0.01, *** p < 0.001, **** p < 0.0001. Data are presented as mean ± SEM from at least three biological replicates.

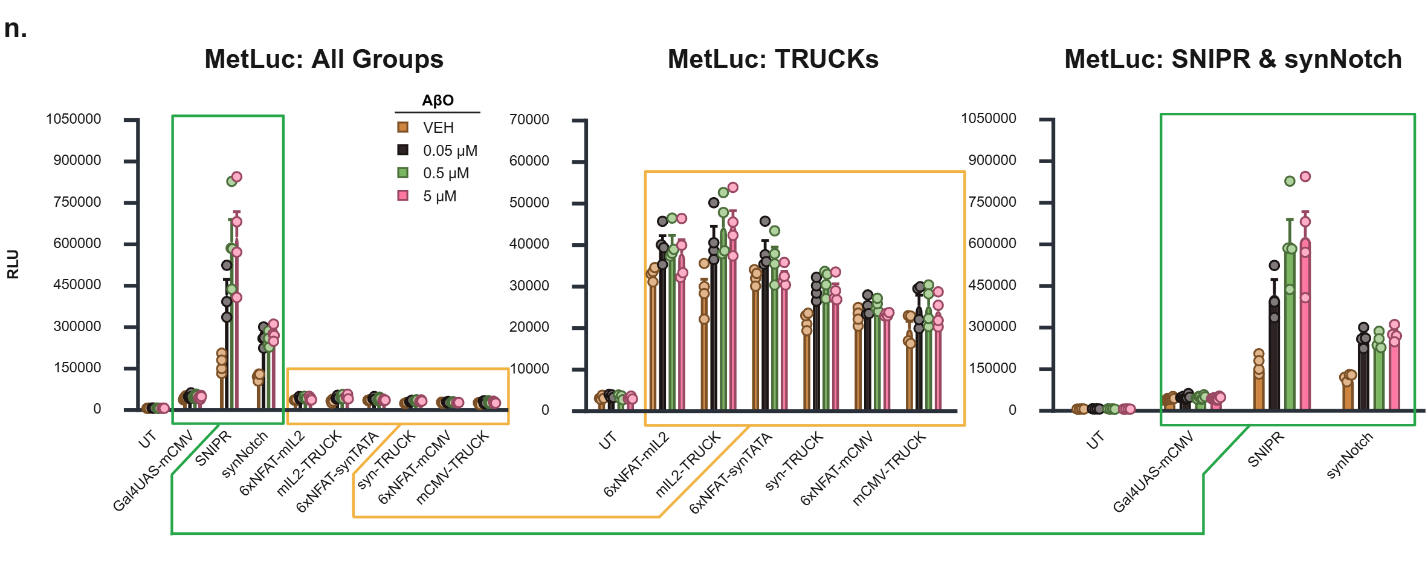

**Supplementary Figure 2n.** Raw MetLuc RLU shown for all groups on a single scale (MetLuc: All Groups, also shown in Fig. 2e) and on separate, scale-optimized plots for TRUCK systems (MetLuc: TRUCKs) and SNIPR/synNotch systems (MetLuc: SNIPR & synNotch) to facilitate visualization. Data are presented as mean ± SEM from at least three biological replicates.

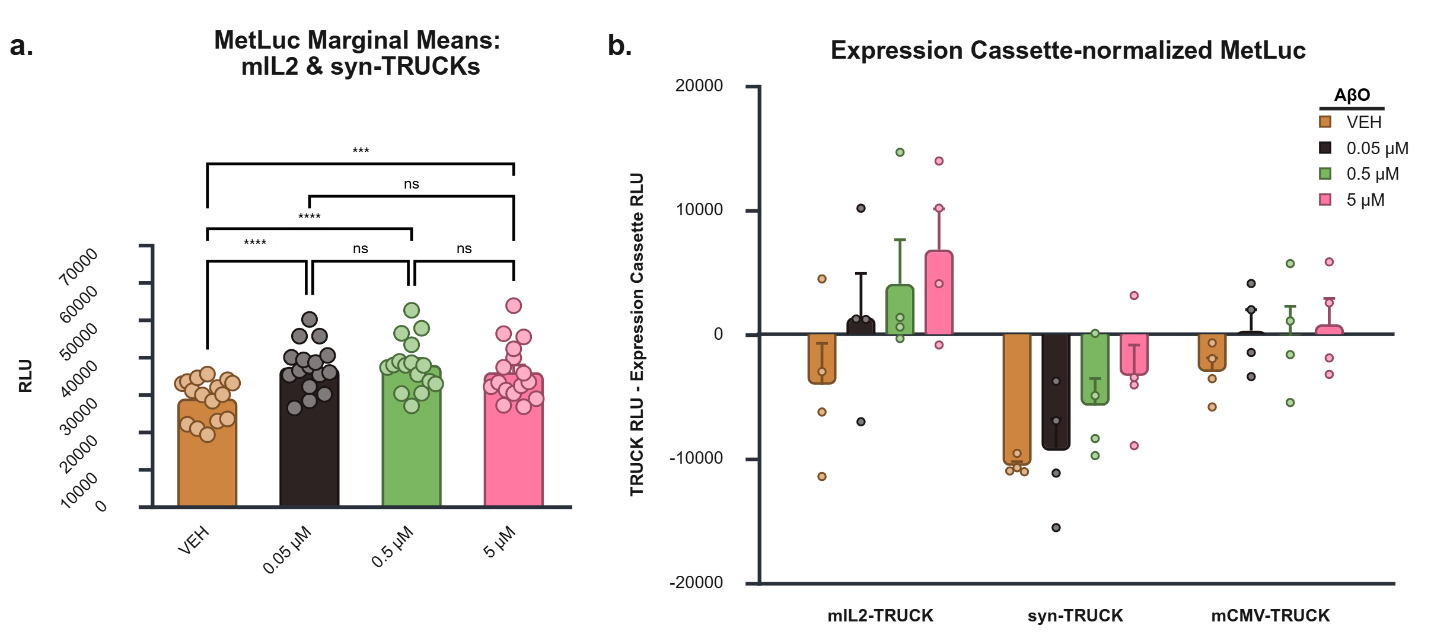

**Supplementary Figures 3a, b**. **a**. Treatment marginal means of MetLuc production across VEH and AβO conditions for mIL2- and syn-TRUCK systems. Treatment marginal means represent the average MetLuc output across all clones within each treatment. Statistical comparisons were performed using Bonferroni-corrected pairwise tests. Additional statistical results are provided in Table S4b. *** p < 0.001, **** p < 0.0001. **b**. MetLuc RLU values for mIL2-, syn-, and mCMV-TRUCK systems normalized to transcriptional leakage from clones expressing only the corresponding 6×NFAT expression cassette (6×NFAT–mIL2, 6×NFAT–synTATA, or 6×NFAT–mCMV). Normalization was performed by subtracting the MetLuc RLU of expression cassette–only clones from the matched full receptor clone for each treatment and replicate. Negative values indicate higher reporter output from expression cassette–only clones than from their corresponding receptor-expressing counterparts. The resulting profiles show negative values under VEH conditions that progressively shift toward positive values with increasing AβO concentration, consistent with partial offsetting of the reduced NFAT-driven output observed in the presence of the Adu-CAR under VEH. Data are presented as mean ± SEM from at least four biological replicates per treatment-by-clone combination.

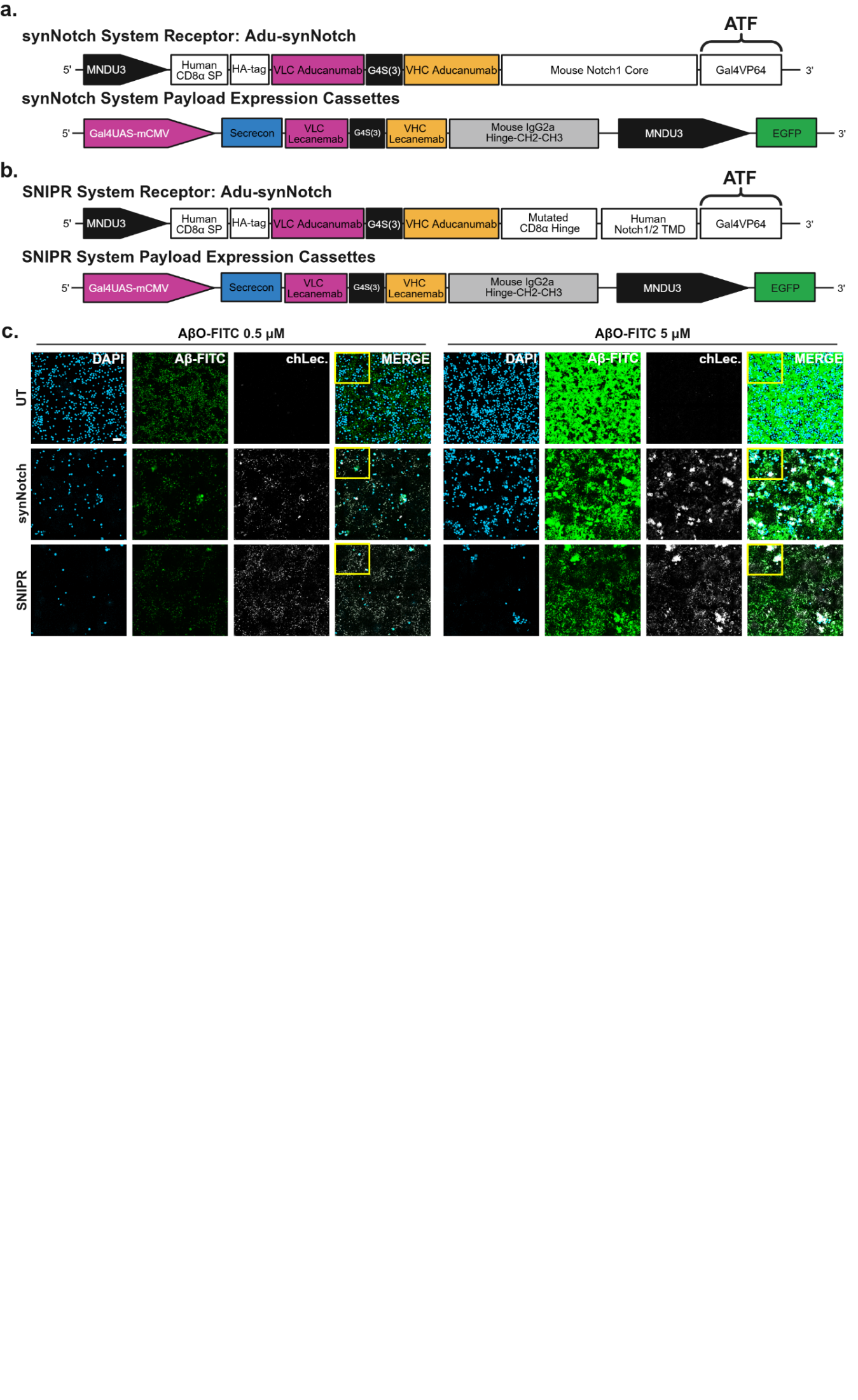

**Supplementary Figures 4a, b.** Architecture of Aβ-targeting (**a**) synNotch and (**b**) SNIPR systems, as described in Sup. Fig. 2a,b, but incorporating an inducible expression cassette encoding chLecanemab only (MetLuc removed). **c**. Representative confocal images of untransduced (UT), synNotch-, and SNIPR-expressing clones under the indicated treatment conditions. Yellow boxes indicate the regions shown in Fig. 4c. Channels: DAPI (nuclei), Aβ–FITC (aggregates), and chLec. (chLecanemab). Images were acquired using a 20× objective (1× zoom). Scale bar, 50 μm.

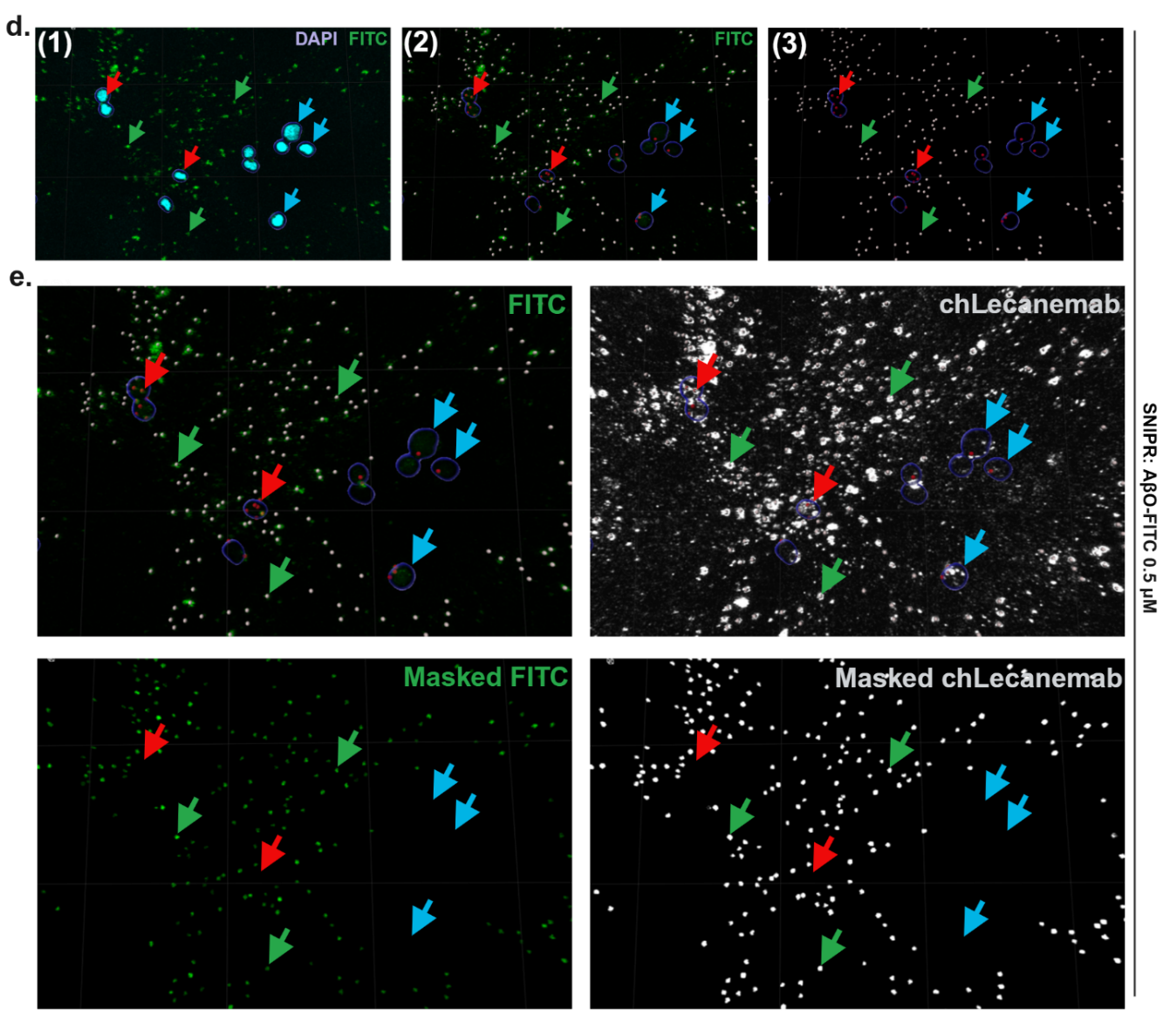

**Supplementary Figures 4d, e. d.** Representative images illustrating the Imaris analysis workflow. (**1**) The Surfaces function was used to generate ROIs around DAPI-positive nuclei (light blue arrows). (**2**) The Spots function was used to generate ROIs corresponding to Aβ–FITC–positive puncta. (**3**) Puncta located within 1 μm of the DAPI-defined nuclear/perinuclear surfaces (red arrows) were excluded from analysis, retaining only puncta outside this region (green arrows) for quantification of Aβ–FITC and chLecanemab voxel intensities. **e**. Images showing the FITC and chLecanemab channels with the 3D ROIs described in (**d**) overlaid. The top row displays the original channels, including voxels both inside and outside the Aβ–FITC–defined puncta used for quantification. The bottom row shows the Masked FITC and Masked chLecanemab channels, in which only voxels within the retained Aβ–FITC–positive puncta (green arrows) are preserved for voxel intensity measurements. Light blue and red arrows indicate voxels outside the retained puncta that were excluded from the analysis.
